# Neutral evolution of snoRNA Host Gene long non-coding RNA affects cell fate control

**DOI:** 10.1101/2023.12.19.572393

**Authors:** Matteo Vietri Rudan, Kalle H. Sipilä, Christina Philippeos, Clarisse Gânier, Victor A. Negri, Fiona M. Watt

## Abstract

A fundamental challenge in molecular biology is to understand how evolving genomes can acquire new functions. Several recent studies have underscored how non-conserved sequences can contribute to organismal diversification in the primate lineage^1–3^. Actively transcribed, non-coding parts of the genome provide a potential platform for the development of new functional sequences^4^, but their biological and evolutionary roles remain largely unexplored. Here we show that a set of neutrally evolving long non-coding RNAs (lncRNA) arising from small nucleolar RNA Host Genes (SNHGs) are highly expressed in skin and dysregulated in inflammatory conditions. SNHGs affect cell fate determination and can behave as evolutionary intermediates to develop new functions^5^. Using SNHG7 and human epidermal keratinocytes as a model, we describe a mechanism by which these lncRNAs can increase self-renewal and inhibit differentiation. SNHG7 lncRNA’s activity has been acquired recently in the primate lineage and depends on a short sequence required for microRNA binding. Taken together, our results highlight the importance of understanding the role of fast-evolving transcripts in normal and diseased epithelia, and inform on how poorly conserved, actively transcribed non-coding sequences can participate in the evolution of genomic functionality.

## Introduction

Advances in the annotation of the genome have yielded the unexpected finding that non-coding sequences are pervasively transcribed^6,7^. While this could imply widespread function^8,9^, some studies have highlighted that the emergence of non-functional and/or redundant sequences is, rather, a by-product of genomic evolution^4,10^.

To distinguish the functional from the non-functional part of the genome the most robust and most commonly used parameter is evolutionary conservation, which in fact is used as the defining characteristic of biological function in its narrower sense^11^. However, use of sequence conservation alone is not sufficient to capture the full extent of genome functionality^12^; recent studies have highlighted how some poorly conserved genomic regions, either regulatory or transcribed, can play important roles in the development of new biological features, or contribute to the phenotypic diversity among species, particularly in the primate lineage^1–3^. Long non-coding RNAs (lncRNAs) are a class of transcripts that generally exhibit a lower degree of sequence conservation when compared to protein-coding or small non-coding RNA. Such conservation can be limited to only a portion of the sequence, to the intron/exon structure or simply to the gene’s position within the genome^13^. In spite of this, several lncRNAs have been ascribed a wide variety of functions through many different mechanisms^14^. While we have accumulated considerable knowledge about the biological roles of a few non-coding RNAs, in most cases their function, if any, remains elusive.

Among the most evolutionarily dynamic tissues is the epidermis, a multi-layered epithelium predominantly composed of keratinocytes that forms the outermost surface of the body. The basal layer contains the epidermal stem cells that maintain the tissue and can either undergo cell division or begin a process of terminal differentiation, during which they migrate towards the surface of the skin, through the spinous and granular layers, and form the protective covering surface of the organism, the cornified layer^15^. Control of the equilibrium between self-renewal and differentiation of keratinocytes is essential for the correct maintenance and repair of the epidermis. Due to its position as the interface with the external environment, the epidermis is a highly evolutionarily dynamic, adaptable tissue, as evidenced, for example, by the relatively high rate of amino-acid substitutions in epidermal proteins, underlying a diverse array of phenotypes between closely related species or even between different human populations^16^. While a few RNA regulators of cell fate have been described in the epidermis^17–19^, the contribution of non-coding parts of the genome to such evolutionary plasticity has received little attention.

Here, we have focussed on small nucleolar RNA Host Genes (SNHGs), a set of highly expressed, extremely poorly conserved lncRNAs whose levels are dysregulated in skin diseases. By using primary human epidermal keratinocytes as a well-established model, we demonstrate the function of SNHGs in regulating the balance between self-renewal and differentiation. Furthermore, comparison of SNHG activity in keratinocytes from multiple mammalian species across different evolutionary distances, provides insights into how new functions can be acquired by the genome.

### SNHGs are a class of poorly conserved, highly expressed long non-coding RNAs that can affect keratinocyte fate

The functional contribution of the major evolutionary conserved signalling pathways in normal and diseased epidermis has undergone extensive investigation. Conversely, the biological role of less conserved transcripts has remained less studied, even though they can be strongly expressed in normal tissue and altered in skin diseases. To identify potentially functional poorly conserved transcripts in the epidermis, we analysed keratinocyte single-cell RNA sequencing (scRNAseq) data^20^. We compared cells from unaffected skin with matched atopic dermatitis (AD) and psoriasis lesions and estimated the conservation of significantly differentially expressed genes using average PhastCons scores from multiple alignments of 100 vertebrate genomes. While many of the affected transcripts were highly conserved coding genes, we were able to identify several very poorly conserved transcripts that were altered in one or both pathological states. As the degree of conservation decreased, we could see an increasing number of lncRNA species, among which we noticed several relatively highly expressed SNHGs (Fig. 1A-B), a class of lncRNA that contain small nucleolar RNAs (snoRNAs) within their introns.

**Fig. 1.**
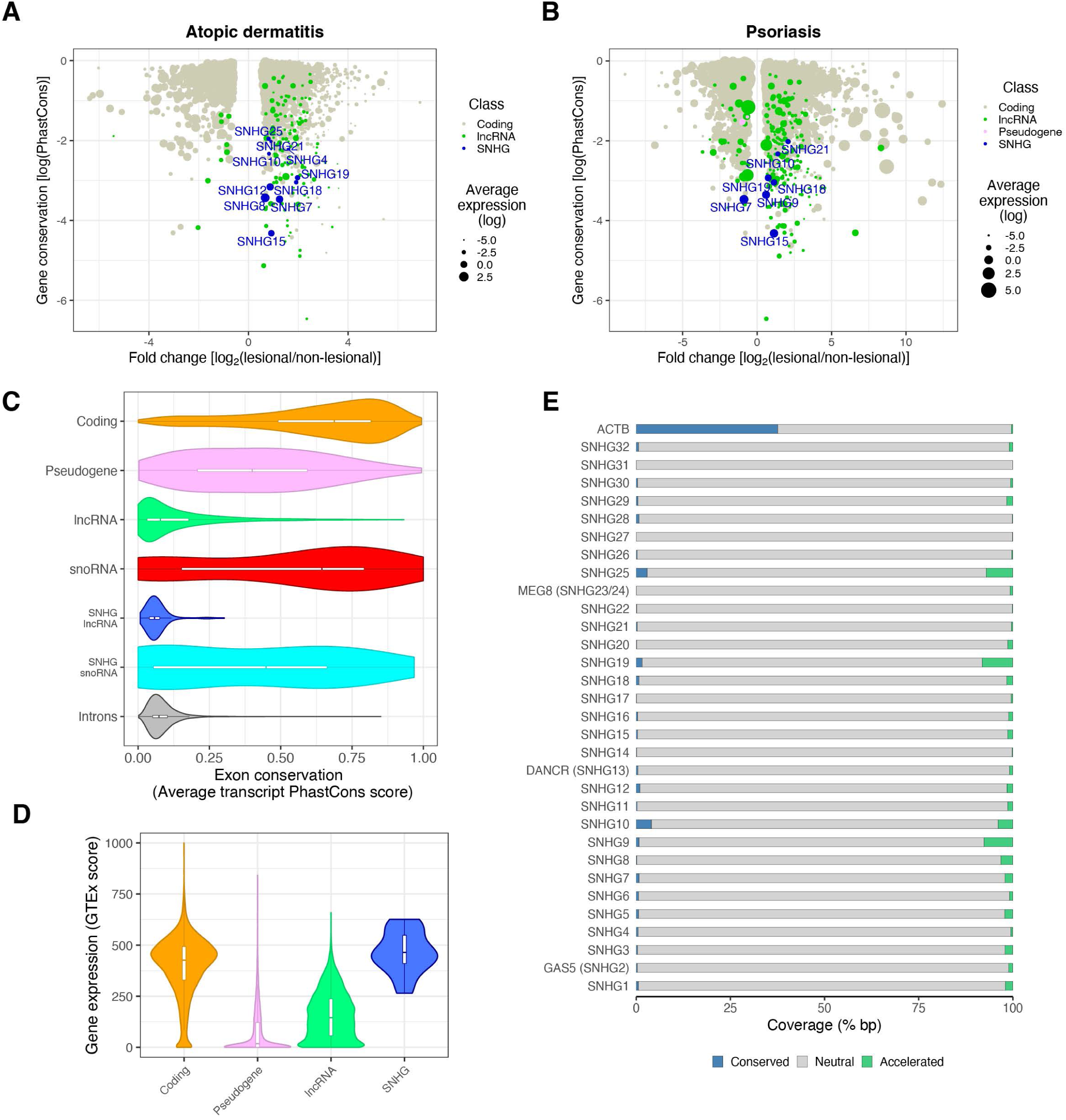
SNHGs encode highly expressed long non-coding RNAs undergoing neutral evolution that are regulated in skin inflammatory conditions. Landscape of differentially expressed genes relative to their conservation in (A) atopic dermatitis and (B) psoriasis. Distributions of (C), exonic sequence conservation and (D), gene expression scores of SNHG lncRNA compared to other classes of transcripts genome-wide. Boxplots within the violins indicate the median and the interquartile range. (E) Rates of evolution of the nucleotides in SNHG exonic sequences assessed by PhyloP scores. Beta-actin (ACTB) is shown as a reference for the rates of evolution in a typical protein-coding gene.

Generally, lncRNAs are expressed at low levels. However, in the case of SNHGs, the production of the abundant intronic snoRNAs requires transcription of the host gene. Indeed, when compared to other transcript classes genome-wide, SNHGs peculiarly display a very low degree of conservation, even relative to known functional lncRNA (Fig S1A) yet retain a level of expression in the same range as most protein-coding genes, as evaluated by using the GTEx scores from data generated by the Gene-Tissue Expression project (Fig. 1C-D, Fig. S1B). SNHG lncRNA sequences contain a relatively high GC content, when compared to other classes of genes (Fig. S1B). SNHG promoters also appear to be less conserved than coding gene promoters (Fig S1C). To control for the possible influence of the phylogenetic scale on our results, we repeated our conservation analyses using average PhastCons scores from multiple alignments of 30 mammals (28 primates) but saw no substantial differences in the outcomes (Fig. S1D-F). We also assessed the rate at which SNHG lncRNAs are evolving. We used PhyloP scores^21^ to gauge what percentages of the exonic sequence of each SNHG are under purifying selection (PhyloP > 2), are evolving neutrally (-2 ≤ PhyloP ≤ 2) or are experiencing an accelerated rate of change (PhyloP < -2). As expected, conserved nucleotides make up a very small fraction of each transcript and while in some cases evidence of accelerated evolution can be observed in up to 8% of base pairs, the vast majority of the sequence of SNHG lncRNAs is evolving neutrally (Fig. 1E). It should be noted that while SNHGs are broadly expressed, there is variation in their levels across tissues, which suggests some degree of regulation of their abundance (Fig. S2).

As dermatitic and psoriatic lesions are characterised by an imbalance balance between self-renewal and differentiation of epidermal cells, we sought to understand if SNHGs could be involved in the regulation of keratinocyte differentiation in normal tissue. We clustered scRNAseq data of keratinocytes from healthy skin donors ^20^ into different states corresponding to stages of differentiation, as described previously^22,23^ (Fig. 2A). We found 13 SNHGs expressed at various levels at multiple differentiation stages (Fig. 2B). We selected the five most highly expressed keratinocyte SNHGs whose expression was also significantly changed in AD or psoriasis (SNHG7, 8, 12, 15, and 19, Fig S3) for further investigation. Expression of these SNHGs could be detected across all clusters, although some enrichment was seen in keratinocytes transitioning from basal to more differentiated cell states, with different SNHGs having maximal expression in either of the two transition states or in certain spinous clusters (Fig. 2C).

**Fig. 2.**
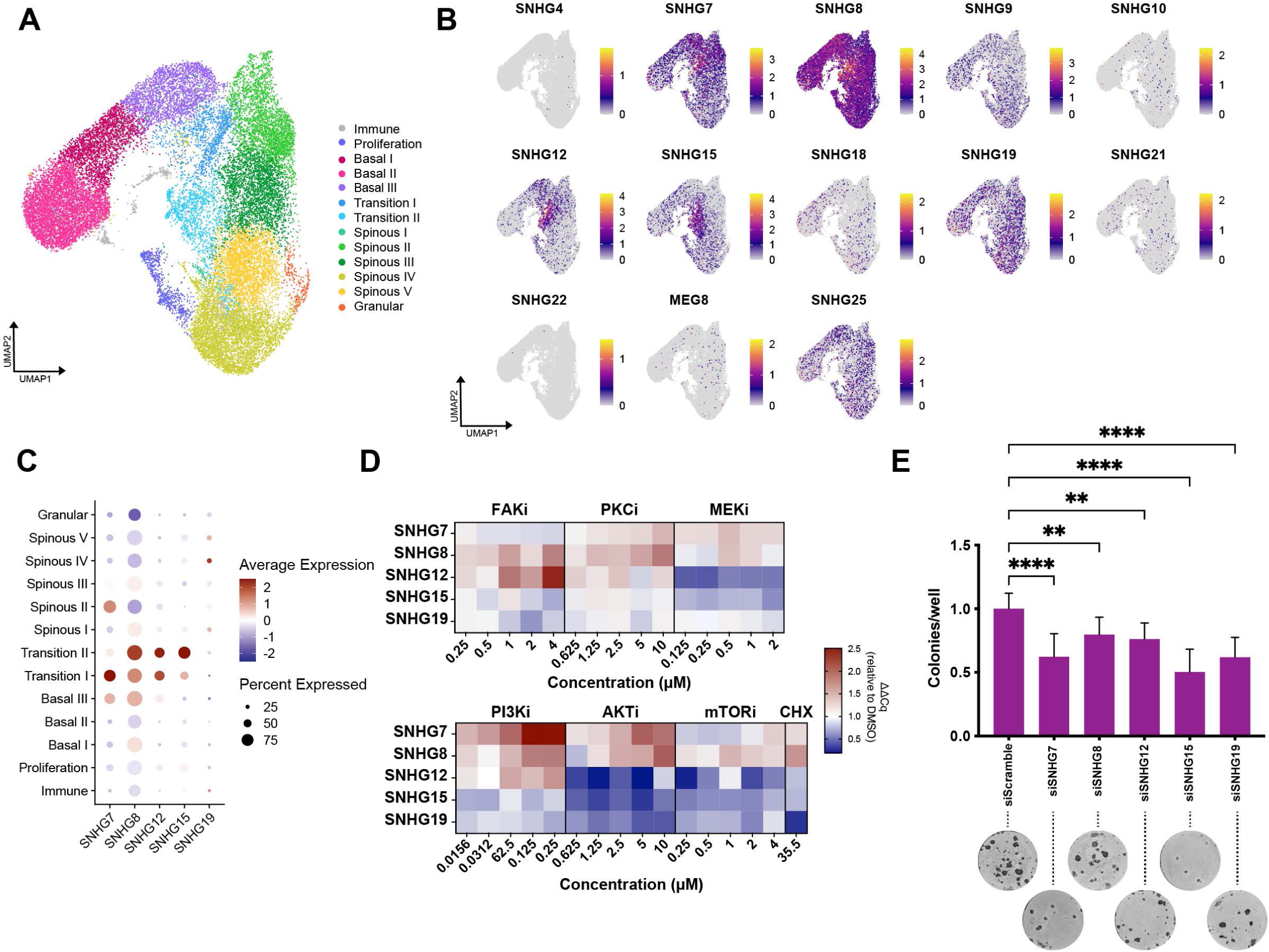
SNHGs expression is regulated in epidermal keratinocytes and can impact their fate. (**A**) UMAP clustering of single-cell RNA sequencing of epidermal keratinocytes. Clusters are shown in different colours and labelled based on marker gene expression. Adapted from Negri et al., 202323. (**B)** Single-cell level expression of the SNHGs detectable in the dataset overlayed on the UMAP clusters. (**C**) Average expression and percentage of keratinocytes expressing five selected SNHGs among the clusters. (**D**) SNHG expression in response to inhibition of multiple signalling pathways. For each inhibitor the change in expression of five epidermally expressed SNHGs compared to DMSO control is shown after 4 h incubation with increasing concentrations of the inhibitor. (**E**) Clonogenicity assay of keratinocytes treated with either non-targeting control siRNA (siScramble) or siRNAs targeting selected epidermally expressed SNHG ncRNA. Data shown in are mean +/- SD. Dunnet’s multiple comparisons test, n = 12 wells. **** p < 0.0001, ** p < 0.01, * p < 0.05.

A variety of signalling pathways participate in regulating the balance between self-renewal and differentiation of keratinocytes^24–28^. The variation in expression among SNHGs during differentiation suggests that they might be under the control of different pathways. Indeed, treatment with a panel of pathway inhibitors had differential effects on the expression of each SNHG as quantified by qPCR. Inhibition of the phosphoinositide-3-kinase (PI3K)/RACα serine/threonine-protein kinase (Akt) pathway increased the levels of SNHG7 and SNHG8, and the latter was also significantly increased after Protein Kinase C or translation inhibition. SNHG12 levels increased upon Focal Adhesion Kinase inhibition, but decreased when the mitogen-activated kinase MEK was blocked. SNHG15 levels decreased in response to Akt or MEK inhibition, while levels of SNHG19 were reduced when the cells were treated with inhibitors of Akt or translation (Fig. 2D, Fig. S4).

In light of these results, we tested whether the presence of SNHG transcripts might have any functional relevance. We used siRNA to knock down each of five highly expressed SNHGs in primary human keratinocytes and performed a clonogenicity assay as a readout of their self-renewal potential^27,29^. Remarkably, a reduction in the expression levels of any of the SNHGs caused a significant decrease in clonogenicity (Fig. 2E; Fig. S5), indicating that neutrally evolving non-coding transcripts arising from snoRNA host genes can affect the balance between self-renewal and proliferation.

### SNHG7 lncRNA promotes self-renewal and prevents differentiation of keratinocytes

Next, we focused on SNHG7 since it is highly expressed in the epidermis (Fig. 2B-C, Fig. S2), its expression is significantly changed in both AD and psoriasis (Fig 1A-B), it strongly regulates clonogenicity (Fig. 2E), and its gene structure is relatively simple: SNHG7 encodes only two alternative lncRNA^30,31^, each about ∼1 kb, and contains two H/ACA box snoRNA genes, SNORA17A and SNORA17B, within its introns (Fig. 3A)^26,27^.

**Fig. 3.**
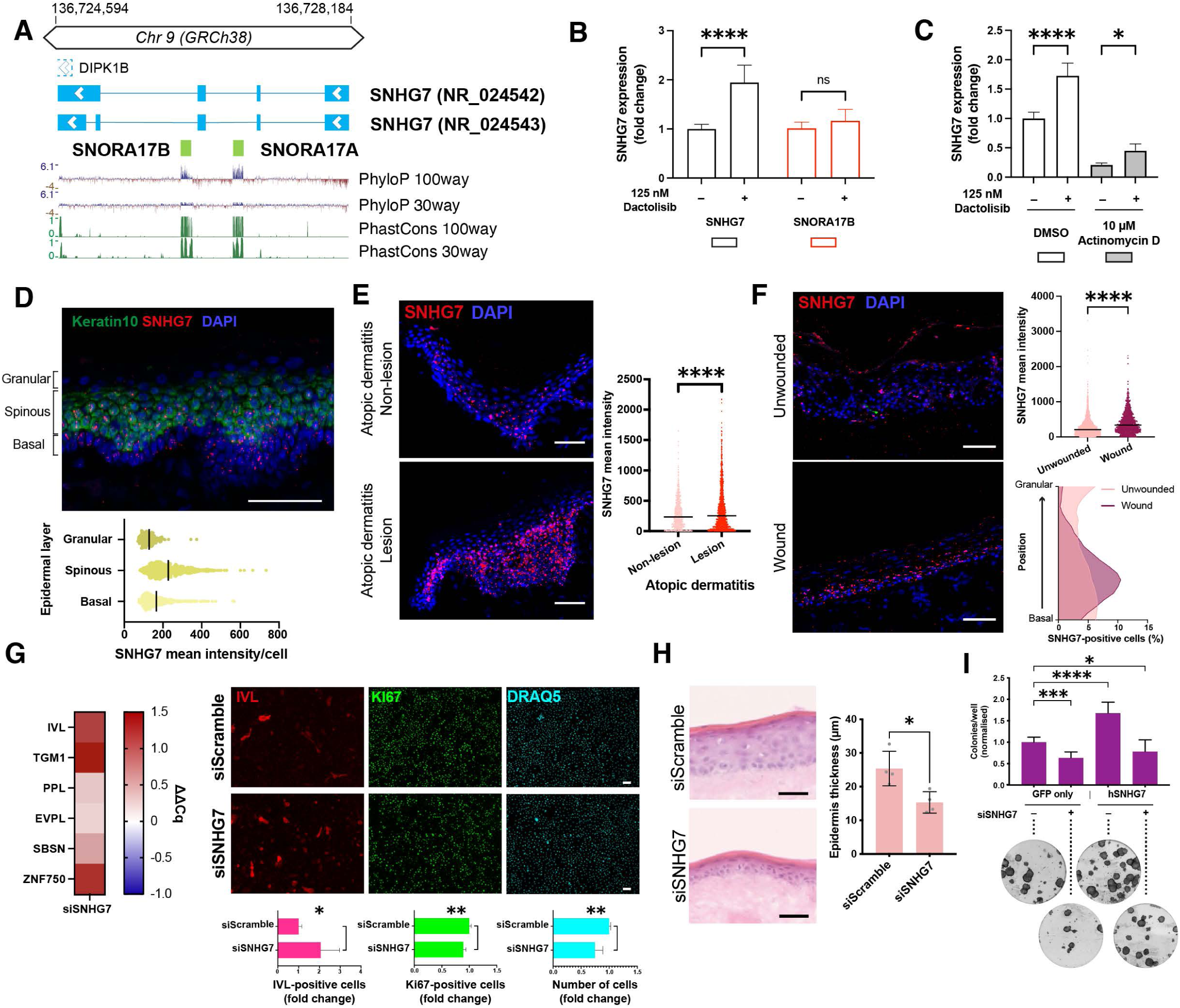
Epidermal SNHG7 lncRNA expression is modulated in homeostatic and non-homeostatic conditions and is necessary to maintain proliferation and prevent differentiation. (**A**) Schematic representation of the SNHG7 genomic locus. Thick lines represent exons, while the thinner lines joining them represent introns. Arrows indicate the direction of transcription. An overlapping 5’UTR of the neighbouring DIPK1B gene is shown as a dashed outline. Conservation tracks are displayed below the schematic. (**B**) Expression of SNHG7 and SNORA17B after 4 h treatment with PI3K inhibitor Dactolisib. n = 9 independent treatments. (**C**) SNHG7 expression after 4 h PI3K inhibition with or without prior 1 h treatment with transcription inhibitor Actinomycin D. n = 5 independent treatments. (**D**) smRNA fish of SNHG7 lncRNA in healthy human facial skin. Keratin 10 RNA staining was used to mark the spinous layer and the expression level of SNHG7 was quantified in each epidermal layer. (**E**) smRNA fish of SNHG7 lncRNA in sections of lesional and non-lesional atopic dermatitis skin, with expression quantification. n > 600 cells. (**F**) smRNA fish of SNHG7 lncRNA in re-epithelialised wounded biopsies. Shown are sections of the areas outside the wound (‘Unwounded’) as well as the newly epithelialised region (‘Wound’). Plots on the right show the quantification of the expression levels in the different areas (top) and the distribution of the SNHG7-espressing cells along the epidermal thickness (bottom). n > 1700 cells. (**G**) Effect of SNHG7 knockdown on keratinocyte differentiation and proliferation. Left heatmap, expression change of multiple differentiation markers in knocked down cells 72 h post-transfection. Right panels, staining of differentiation (IVL) and proliferation (Ki67) markers 96 h post-transfection with quantifications. n ≥ 4 independent transfections. (**H**) Effect of SNHG7 knockdown on the ability of keratinocytes to form a new epithelium. n = 4 DEDs. (**I**), Rescue of the SNHG7 knockdown phenotype by lncRNA overexpression. Clonogenicity assay quantification with representative images. Scale bars, 50 µm. Data shown in all bar plots are mean +/- SD. Lines in dot plots indicate the median. (**B**), (**E**), (**F**), (**G**) and (**H**) Two-tailed unpaired t-test. (**C**) Šìdàk’s multiple comparisons test. (I), Dunnet’s multiple comparison test. **** p < 0.0001, ** p < 0.01, * p < 0.05.

The modulation of SNHG7 lncRNA transcription is linked to the expression of the snoRNAs hosted in its introns. Our screen of pathway inhibitors showed that SNHG7 was markedly upregulated upon PI3K inhibition (Fig. 2E, Fig. S4). Intriguingly, this was not mirrored by the levels of a snoRNA hosted by SNHG7 (Fig. 3B). When we inhibited transcription with Actinomycin D before blocking PI3K, a significant increase of SNHG7 could still be detected, while the transcription-dependent, PI3K-mediated up-regulation of ERBB3^32^ was completely prevented (Fig. 3C, Fig. S6A). These results point to the presence of a post-transcriptional regulatory mechanism for the lncRNA, allowing its expression to be decoupled from the snoRNA.

In vivo, SNHG7 was present in scattered cells throughout the living layers of human epidermis, with higher expression in the basal and spinous layers (Fig. 3D) as shown by single-molecule RNA fluorescence *in-situ* hybridisation (smRNA FISH), and in agreement with our scRNAseq analysis. In non-homeostatic conditions, we observed alterations in SNHG7 lncRNA levels. SmRNA FISH of AD lesions showed an increase in SNHG7 expression (Fig. 3E), which was consistent with scRNAseq from AD patients.

To experimentally demonstrate the modulation of SNHG7 in non-homeostatic conditions, we used *ex vivo* wounding of human skin explants. The re-epithelialising keratinocytes exhibited increased levels of SNHG7 lncRNA, including more prominent expression in basal cells (Fig. 3F). This supports the idea that SNHG7 participates in the regulation of the balance between self-renewal and differentiation in the context of epidermal repair. Our results also indicate that keratinocyte SNHG7 expression can increase in response to perturbation of homeostasis in the absence of circulating immune cells.

To further characterise the reduction in clonogenicity observed upon down-regulation of SNHG7 we stained keratinocytes with antibodies to the proliferation marker Ki-67 and the differentiation marker Involucrin (IVL). We observed a reduced rate of proliferation and an increased percentage of differentiated cells upon SNHG7 knockdown. The induction of differentiation was confirmed by the increase in mRNA levels of a range of differentiation markers (Fig. 3G). Knockdown cells also displayed a reduced ability to form an epidermis on de-epidermised dermal substrates (Fig. 3H). We did not detect marked alterations in the distribution of epidermal markers in the reconstituted tissues (Fig. S6B). Off-target effects were excluded by deconvolution of the siRNA pools used in our experiments (Fig. S6C-D). The effects of SNHG7 knockdown were not due to induction of apoptosis since there was no increase in the number of cells expressing cleaved caspase 3 (Fig. S6E).

We also ruled out any potential contribution of the snoRNA genes hosted within SNHG7 to the knockdown phenotype. Firstly, no reduction in protein translation as assessed by the OP-Puro assay was seen in knockdown cells (Fig. S6F). Secondly, consistent with the post-transcriptional nature of RNA interference, siRNA-mediated knockdown of SNHG7 only affected the exonic portion of the gene (Fig. S6G). We further demonstrated that the effects we observed were due to the deficiency in SNHG7 lncRNA by overexpressing its spliced form (NR_024542). Since the siRNA is able to target both the endogenous and the exogenous lncRNA we first identified the minimum siRNA concentration necessary to see a clear phenotype (Fig. S6H). Overexpression of SNHG7 lncRNA led to a nearly complete rescue of the knockdown phenotype (Fig. 3I, Fig. S6I-K).

Overexpression of SNHG7 caused a significant increase in clonogenic potential (Fig. 3I). Additionally, stable overexpression of SNHG7 allowed the keratinocytes to be kept in culture for seven additional passages before losing their self-renewal capacity (Fig S6L). This, together with the high levels of expression of SNHG7 during keratinocyte commitment and early differentiation (Fig. 2C), prompted us to investigate whether SNHG7 might affect suspension-induced differentiation of disaggregated human keratinocytes^27^. While we did observe a more rapid induction of the pro-commitment phosphatase DUSP6 upon SNHG7 knockdown (Fig. S6M), the kinetics of differentiation in suspension were largely unchanged. In addition, overexpression of SNHG7 did not prevent the loss of colony forming ability with time in suspension (Fig. S6N). These results indicate that SNHG7 influences the propensity of keratinocytes to undergo self-renewal, but its activity can be overridden by the strong differentiation stimulus of detachment from the extracellular matrix.

### SNHG7 function has evolved recently in the primate lineage

The low sequence conservation of SNHGs raises the possibility that their effects on the regulation of cellular fate can be acquired over relatively short evolutionary timescales. However, cases have been described of lncRNA which maintained their function across large evolutionary distances, in the absence of widespread sequence conservation^33^. In order to distinguish between these alternative scenarios, we assessed the biological activity of SNHG7 in night monkey (*Aotus trivirgatus*) and mouse (*Mus musculus*) primary keratinocytes. Given the absence of genomic sequence information for *A. trivirgatus*, we first performed 5’ and 3’ Rapid Amplification of cDNA ends (RACE) followed by sequencing of the SNHG7 lncRNA isolated from the night monkey cells. We could identify at least two transcripts (Supplementary Data 1), both of which displayed relatively high sequence similarity to the human lncRNA in their 3’ region (Fig. 4A; Supplementary Data 2-3). Conversely, the murine Snhg7 sequence retains little homology to the human lncRNA throughout its sequence (Fig. 4A; Supplementary Data 4). Inhibition of PI3K increased the levels of SNHG7 in night monkey cells, as in human, while it did not affect its expression in mouse keratinocytes (Fig. 4B).

**Fig. 4.**
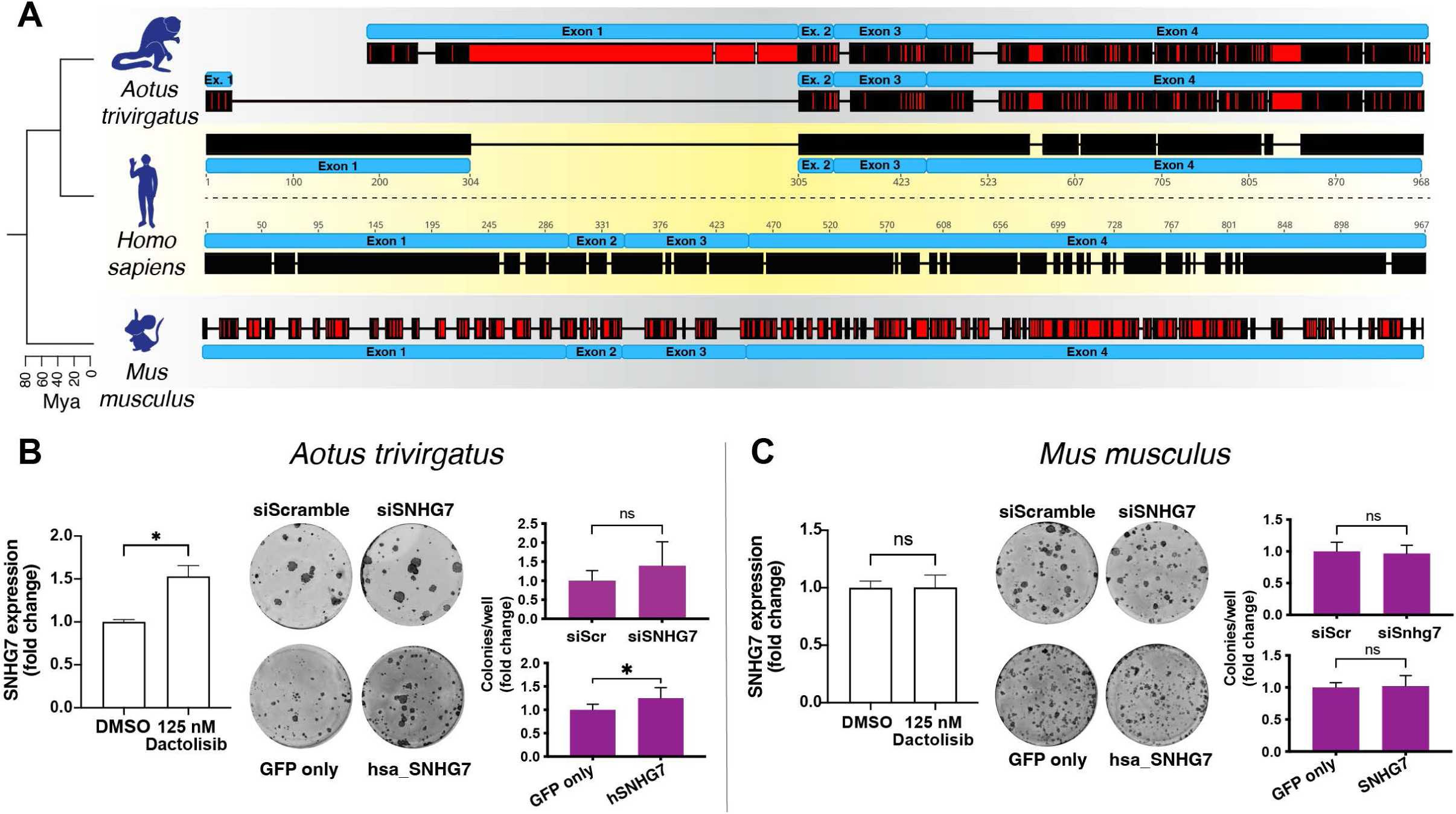
Human SNHG7 lncRNA can affect cell fate determination in primate species where its function is lost. (**A**) Alignment of human SNHG7 lncRNA sequence with its orthologs in South American night monkey (Aotus trivirgatus, top) and mouse (Mus musculus, bottom). A cladogram on the left shows the evolutionary relationship among the species with approximate time separating them. A black colour represents a more conserved sequence, while a red colour represents sequence divergence, thin lines represent gaps in the alignment. (**B**) and (**C**) Regulation and activity of SNHG7 in night monkey (B) and mouse (C) keratinocytes. For each species, shown are expression of SNHG7 after 2 h PI3K treatment (left panels) and clonogenicity assays following either downregulation of endogenous SNHG7 or ectopic overexpression of human SNHG7 with quantifications (right panels). Data shown are mean +/- SD. Unpaired two-tailed t-tests, n ≥ 3 independent treatments or n ≥ 6 wells for clonogenicity assays. * p < 0.05

Although SNHG7 expression in night monkey cells was regulated by PI3K, the colony formation ability of night monkey and mouse keratinocytes was not affected by SNHG7 knockdown (Fig. 4B; Fig. S7A-B). This indicates a rapid acquisition of functionality to accompany the variation in sequence. We therefore sought to investigate whether human SNHG7 was sufficient to affect the proliferation/differentiation balance in non-human keratinocytes. To this end, we stably overexpressed the human SNHG7 lncRNA in night monkey and mouse keratinocytes. Human SNHG7 lncRNA was able to increase the colony forming capacity of night monkey keratinocytes but had no effect on the mouse cells (Fig. 4B; Fig. S7B-C).

Our data therefore points to the presence of elements in the human SNHG7 sequence that are able to influence cell fate, whose efficacy however appears conditional to the presence of additional factors and/or conditions within their cellular environment.

### Presence of microRNA response elements affects SNHG7 activity

Next, we sought to characterise the molecular mechanism of SNHG7’s action. RNAseq of primary human keratinocytes at 24 and 48 h after SNHG7 knockdown (Fig. S8A) revealed that the differentially regulated genes were predominantly involved in control of proliferation, cell motility and cell adhesion (Fig. 5A-B; Supplementary Tables 1-2). Already 24 h after transfection, the differentially expressed genes exhibited a notable bias towards downregulated genes (Figure 5A), which is consistent with the idea that SNHG7 might be a miRNA regulator, as reported in cancer settings^34^. Importantly, the changes in gene expression occurred before cells upregulated differentiation-associated genes (Fig. S8B).

**Fig. 5.**
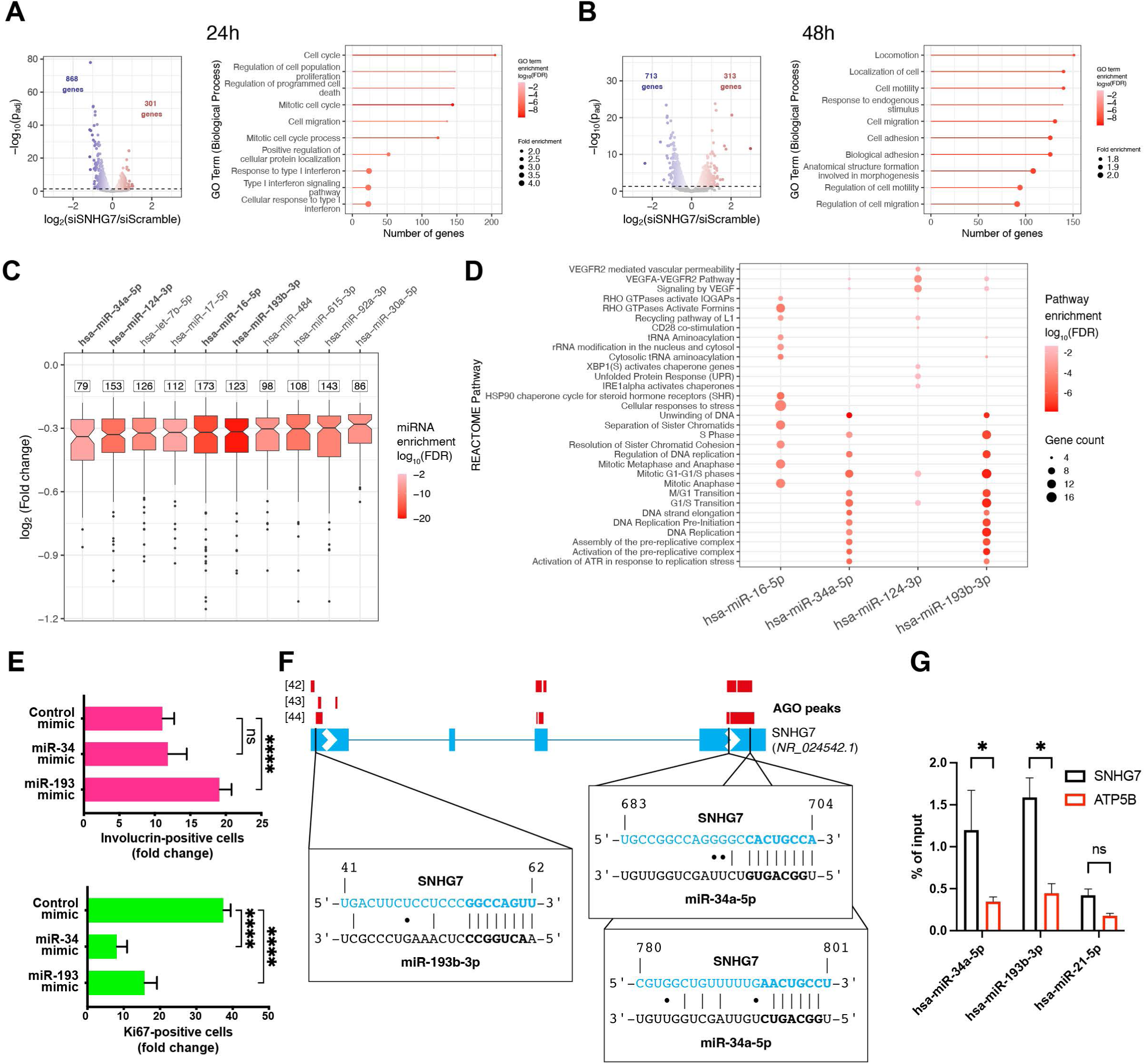
SNHG7 lncRNA-regulated genes are enriched in targets of SNHG7-interacting miRNAs. (**A**) and (**B**) Volcano plot (left) and GO-term enrichment analysis (right) of significantly (p_adj_ < 0.05) differentially expressed genes 24 h (**A**) or 48 h (**B**) after siSNHG7 transfection. The colour of the lollipops indicates the significance of the GO term enrichment. (**C**) miRNA response element enrichment in significantly (p_adj_ < 0.05) downregulated genes 24 h post-transfection. The number of target genes for each miRNA is shown above the box plot of their differential expression. miRNAs are sorted based on the median downregulation of their target genes. Candidate miRNAs with MREs in the SNHG7 sequence are highlighted in bold. The colour of the boxes indicates the significance of the enrichment. (**D**) Functional enrichment of the downregulated targets of candidate miRNAs against the REACTOME pathway database. The colour of the points indicates the significance of the enrichment. (**E**) Effect of miR-34a-5p and mir-193-3p mimics on keratinocyte proliferation and differentiation. (**F**) Schematic of miRNA binding site location on SNHG7 with location of Ago1/2 peaks from published Clip datasets. Thick lines represent exons, while the thinner lines joining them represent introns. Arrows indicate the direction of transcription. Vertical bars represent Watson-Crick pairings. Seed sequences on miRNAs and MREs on SNHG7 are highlighted in bold. (**G**) Enrichment of SNHG7 after pulldown of biotinylated candidate miRNAs, a non-biotinylated scramble control or a biotinylated miRNA without MREs in SNHG7 (miR-21-5p). Enrichment for a control gene without MREs for any of the miRNAs (ATP5B) is also shown. n = 2 pulldown experiments. Fisher’s LSD test. Data shown are mean +/- SD. * p < 0.05

Analysis of the intracellular distribution of SNHG7 showed a predominantly cytosolic location, similar to mRNAs and compatible with the ability to bind miRNAs (Fig. S8C). To identify which microRNAs could be involved in the biological activity of SNHG7, we uploaded the list of significantly downregulated (p_adj_ < 0.05) genes 24 h after transfection into MIENTURNET^35^ and scanned for significant enrichment of miRNA targets based on experimentally validated miRNA-target interactions from MiRTarBase (Fig. 5C; Supplementary Table 3). The same analysis was also performed for genes significantly downregulated after 48 h (Fig. S8D; Supplementary Table 3). Among the ten most significantly enriched miRNAs, six have the potential to bind to SNHG7. miR-34a-5p and miR-193b-3p have “canonical” miRNA-response elements (MRE)^36^ within the sequence of SNHG7, miR-124-3p can form a non-canonical type of binding that has been previously reported^37^, miR-16-5p, miR-484 and miR-615-3p can potentially form 6-mer “marginal” pairing with SNHG7. Although this latter type of binding is less likely to be conducive to regulation^37^ miR-16-5p targeted the highest number of downregulated genes at both 24 h and 48 h post transfection.

To further investigate the interplay between SNHG7 and miR-16-5p, miR-34-5p, miR-124-3p and miR-193-3p, we performed pathway enrichment analysis of significantly downregulated genes targeted by each miRNA against the REACTOME database at 24 h post transfection (Fig. 5D; Supplementary Table 4). miR-34a-5p and miR-193b-3p targets were predominantly associated with cell cycle and proliferation, while miR-16-5p and miR-124-3p were more associated with signalling and cellular stress. Since the SNHG7 knockdown phenotype is related to the balance between proliferation and differentiation, we further narrowed down our pool of candidates to miR-34a-5p and miR-193b-3p. The experimentally validated targets of these two miRNAs display a strong bias towards downregulation in the siSNHG7 RNAseq data, unlike targets for miR-21-5p which were not enriched among the significantly downregulated genes and does not have an MRE within the sequence of SNHG7 (Fig. S8E). Similarly, looking at the distribution of all predicted targets for these two miRNA families confirmed a significant shift towards downregulation for both miRNAs, which was not observed in the case of miR-21-5p (Figure S8F). SmRNA *in situ* hybridisation showed that both of these miRNAs are expressed in keratinocytes (Fig. S8G). While the activity of the miR-34-5p family of miRNA in epidermal cells has been described^38^, the effect of miR-193-3p on keratinocytes is not known. Consistent with the SNHG7 phenotype, transfection of either miR-34a-5p or miR-193b-3p caused a marked reduction in keratinocyte proliferation, accompanied by a significant increase in differentiation in the case of the miR-193b-3p mimic (Fig. 5E).

In support of the potential for miRNA-based regulation, analysis of published Ago-CLIP datasets^39–41^ showed the presence of peaks of Argonaute binding on the SNHG7 lncRNA transcript overlapping with the MREs for miR-193-3p and miR-34-5p (Fig. 5F). To verify the binding between SNHG7 and the miRNAs, we transfected the cells with biotinylated versions of miR-193b-3p and miR-34a-5p and a control miRNA for which no MRE is present on SNHG7 (miR-21-5p), then performed streptavidin mediated pulldown followed by qPCR. Both miRNAs, but not the negative controls, were able to co-precipitate SNHG7 at higher levels than a negative control gene (ATP5B) and compatible with positive control target genes (Fig. 5G; Fig. S8H).

We also checked whether the miRNA levels increased after SNHG7 was downregulated, as this could indicate that SNHG7 could operate via Target-Directed miRNA Degradation (TDMD) by SNHG7^42,43^. However, we observed no consistent increase of either miR-34a-5p or miR-193b-3p 48 h after transfection (Fig. S8I).

Next, we sought to confirm whether either miRNA could contribute to the biological activity of SNHG7. The night monkey SNHG7 does not contain MREs for either of the candidate miRNA families. We thus used night monkey keratinocytes as a null background to assess the activity of human SNHG7 carrying mutations in either the miR-193-3p or miR-34-5p MREs, as well as a control mutation of similar size in the central region of the transcript (Fig. 6A; Fig. S9A). Stable transfection of miR-193 or control mutant human SNHG7 in night monkey keratinocytes increased their clonogenic potential similar to the wild-type human transcript. However, mutation of the miR-34 MREs abolished the activity of the exogenous transcript (Fig. 6B; Fig. S9B).

**Fig. 6.**
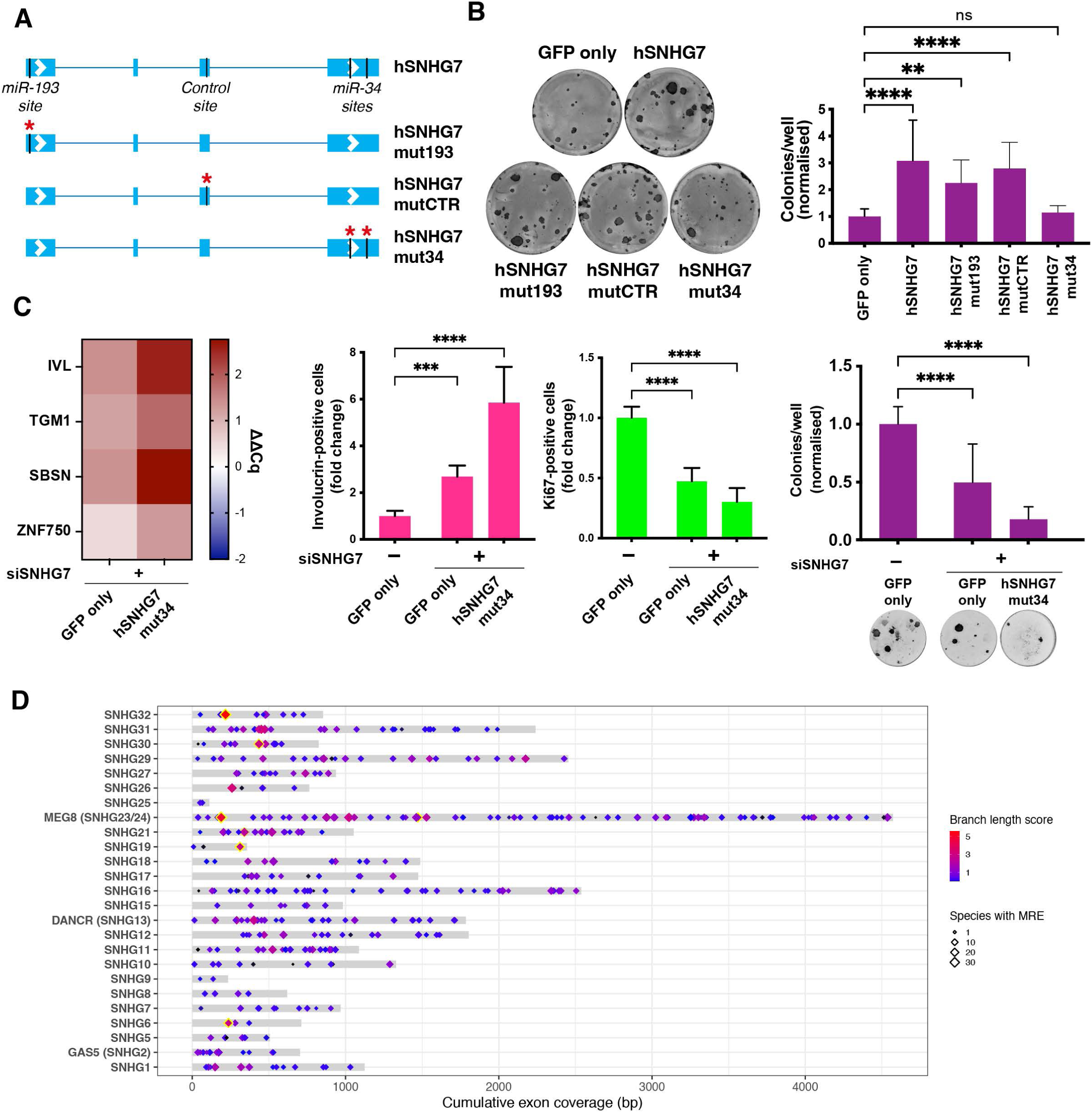
SNHG7 activity requires the presence of miR-34 binding sites. (**A**) Schematic of SNHG7 mutants. (**B**) Effect of the overexpression of human SNHG7 mutants on the clonogenicity of night monkey keratinocytes. (**C**) Rescue of the SNHG7 knockdown phenotype by overexpression of SNHG7 deficient for miR-34 binding sequences. Left heatmap, expression change of multiple differentiation markers in knocked down cells 72 h post-transfection. Middle barplots, staining of differentiation (IVL) and proliferation (Ki67) markers 96 h post-transfection with quantifications, n ≥ 10 independent transfections. Right, clonogenicity assay, n ≥ 18 wells. Data shown are mean +/- SD. (**D**) Distribution and conservation of MREs for deeply conserved miRNAs in SNHGs. Yellow borders mark MREs that pass the TargetScan conservation threshold. (**B**), (**C**) Dunnet’s multiple comparisons tests. **** p < 0.0001, *** p < 0.001, ** p < 0.01, * p < 0.05

Expression of the wild-type SNHG7 lncRNA was able to rescue the knockdown phenotype in human keratinocytes (Fig. 3I). Conversely, expression of the miR-34 MRE mutant was unable to produce any attenuation of the effects of SNHG7 knockdown in human keratinocytes (Fig. 6C; Fig. S9C), further indicating a role for the miR-34-5p MRE in SNHG7 biological activity. Our results thus indicate that SNHG7 regulates clonogenicity at least partially through a miR-34-5p response element and therefore that rapidly evolving, short elements in highly expressed non-coding RNA can confer or contribute to the acquisition of new functions over short evolutionary distances.

Short elements such as MREs can potentially arise and disappear rapidly during evolution. In fact, assuming a uniform base frequency and in the absence of selection, a MRE matching any deeply conserved miRNA could potentially occur by chance every ∼150 bp. In the case of SNHG7, the functional MRE has emerged recently. However, if other SNHGs could function in this way, one would expect to see a range of sequence conservation among MREs in their sequence. To evaluate this, we used the TargetScan 7 pipeline to identify and score the conservation of MREs in SNHG lncRNA sequences (Supplementary Table 5). For this analysis, we only considered broadly conserved miRNA families and only 7-mer and 8-mer MREs. Additionally, we excluded SNHGs that had significant overlap with protein-coding genes and transcripts that had exons overlapping the snoRNAs, as this would introduce a conservation bias. The majority of the sites we identified were shared only among primates, although about a quarter were conserved more deeply (Fig. S9D) and a few passed the TargetScan conservation thresholds^44^ (Fig. 6D). A comparison of MRE conservation between the 25 SNHGs included in the analysis and a sample of 250 3’ untranslated regions (UTRs) of coding genes shows similar distributions, albeit with a higher number of deeply conserved sites in UTRs (Fig. S9E). This increased presence of strongly conserved MREs in UTRs could be reflective of their common susceptibility to miRNA regulation (whereas only a subset of SNHGs might be affected by miRNAs), and/or of an increased difficulty in identifying SNHG orthologs in more distant species. Altogehter, these data suggest that MREs in SNHGs can in some case become selected.

## Discussion

In this study, we have identified a group of SNHGs, a class of highly expressed, poorly conserved non-coding genes, that have a pronounced effect on cell fate decisions in human keratinocytes. SNHGs are a known but understudied class of transcripts: they have been reported to possess biological activity in cancer cells^34^, but only relatively few studies have explored their role in adult tissues^17,45^. The epidermal expression of SNHGs can be controlled by specific signalling pathways; their levels vary during the early phases of epidermal keratinocyte differentiation, and they are significantly altered in certain skin inflammatory conditions and during wound healing. Knockdown of several of the SNHGs in primary human keratinocytes led to a strong reduction in self-renewal capability. This result is consistent with an early report on the activity of another SNHG (DANCR) in the epidermis^17^; however, this gene did not appear in the scRNAseq dataset we analysed. Our data thus paints a picture in which the interplay among multiple signalling pathways affects the levels of several neutrally evolving, non-coding transcripts that influence whether a cell undergoes self-renewal or initiates differentiation and potentially also contribute to stress responses or pathological phenotypes.

In the case of SNHG7, the effect on cell fate determination was acquired recently in the primate lineage. Xenotopic expression of the human transcript could moderately increase the clonogenic capacity of night monkey keratinocytes but did not affect mouse epidermal cells. This differential effect on evolutionarily more distantly related species suggests that SNHG7 activity relies on the presence of context-specific *trans*-acting factors that are present in the monkey cells but absent in the mouse. It is also interesting to note how expression of the human transcript in the monkey cells had a similar effect to the overexpression of the transcript in human keratinocytes; however, unlike human, *A. trivirgatus* cells were not dependent on SNHG7 to fully maintain their ability to self-renew in culture conditions. This indicates that highly expressed, neutrally evolving non-coding transcripts can exhibit a remarkably dynamic functional evolution.

The capacity of SNHG7 to influence keratinocyte fate determination was at least partially dependent on the presence of MREs for a specific miRNA family, miR-34-5p. Regulation of gene expression by competition for the binding of miRNAs has been widely described^46^; however, its effectiveness is severely constrained by strict stoichiometric and biochemical parameters^47,48^. Despite this, several instances have been reported where non-coding RNAs of different types can effectively act as sponges for miRNA, mostly in cancer settings^49–51^ but also in the regulation of neuronal activity^52^ and muscle differentiation^53^. miR-34-5p is known to regulate keratinocyte differentiation in mouse^38^. Nevertheless, human SNHG7 was ineffectual in regulating cell fate in cells from this species, suggesting that additional agents or conditions are involved in its activity. It should also be noted that we cannot exclude the potential contribution of other, miRNA-independent, molecular mechanisms to the action of SNHG7. A full dissection of SNHG7-interacting molecules and detailed investigation of the biochemistry involved in its biological activity will be required to fully understand how it operates.

The expression of intronic snoRNAs, which do not possess their own promoters, is dependent on the transcription of the gene within which they reside^54^. This implies that in the case of SNHGs, non-coding transcripts retain some selective pressure to be transcribed because of the presence of the snoRNA. The necessity of their transcription for snoRNA production and the lack of strong sequence constraints deriving from the need to code for protein thus potentially optimally places SNHG lncRNAs to function as substrates for the evolution of new functional genetic elements or structures^4,55^. Indeed, the absence of SNHG7 activity on the self-renewal/differentiation balance in species where its sequence does not include the MRE for miR-34-5p makes it a paradigmatic example of an “evolutionary spandrel”: a transcript whose presence is a by-product of the expression of a separate RNA species (in this case two snoRNAs) and whose sequence is able to neutrally accumulate mutations and act as hotspots for the acquisition of new effects, which can then be subject to the action of drift or selection^5,11,55^.

The reliance of the biological effect of SNHG lncRNAs on the presence of short MREs together with their remarkably low sequence conservation suggests that the fine-tuning of cell fate decisions and potentially other cellular phenotypes could be influenced by the rapid emergence of these genetic elements in abundant transcripts lacking strong constraints on sequence conservation. Due to the short length of MREs, the emergence of sequences with the potential to confer new functions is likely to occur relatively frequently. Most of the MREs, however, would probably be non-functional because (1) not all SNHGs are likely to operate through the same molecular mechanism, and (2) the additional limitations imposed by the context-dependent factors that limit the effectiveness of miRNA sequestration are likely to constrain the intensity and the spread of the effects of SNHGs. We speculate that upon emergence these effects are unlikely to be very strong or ubiquitous, because an excessively penetrant new property would probably be selected against. This is in fact corroborated by the noticeable but relatively limited magnitude of the effect that human SNHG7 expression has on night monkey keratinocytes.

Our data are consistent with a model in which new, limited-scope effects can emerge out of highly expressed, poorly conserved transcripts over brief evolutionary periods and contribute, for example, to the fine-tuning of the balance between cell proliferation and differentiation in adult tissues. Our findings experimentally corroborate the constructive neutral evolution theory and highlight the potential role played by neutrally evolving sequences in the evolution of new genomic functions in the primate lineage^4,55,56^. The extent and the impact of this phenomenon on general organismal diversification and adaptation to changing environments remain to be determined and represent fascinating open questions ripe for further exploration.

## Methods

### scRNAseq analysis

Single-cell skin data from Reynolds et al.^20^ was subsetted to include keratinocytes isolated from lesional and non-lesional skin of either atopic dermatitis or psoriasis patients. The two object thus obtained were normalised and scaled using default parameters. The differentially expressed genes between lesional and non-lesional atopic dermatitis and psoriasis skin were identified separately for each condition by running the FindMarkers function with parameters test.use=wilcox and logfc.threshold=log2(1.5) and keeping all genes with an adjusted pvalue < 0.01. Analysis of the healthy skin data was performed as described previously^23^.

### Genome-wide analysis of conservation, expression, and GC content

In order to assess the degree of conservation of different gene classes in the scRNA seq data or genome-wide, the PhastCons scores for all annotated transcripts were calculated by averaging the scores of their exonic portions. Exon coordinates were extracted from the UCSC RefSeq table downloaded from the hg38 human genome assembly on the UCSC Genome browser and used to interrogate the 100-vertebrate (Fig.1A-C, Fig. S1A,D) or 30-mammals (Fig. S1E-G) conservation PhastCons scores (range: 0-1). Transcripts were classified according to the gene_biotype field in the ncbiRefSeqLink table downloaded from the same source. In the genome-wide plot (Fig. 1C, Fig. S1B), exonic scores were averaged for every transcript. Intron coordinates from 10,000 randomly selected transcripts were used to serve as a “neutrally evolving” control. For the promoter conservation analysis, the same process was applied to the 500 bp preceeding the transcription start site.

Genome-wide expression scores were obtained from the gtexGeneV8 table downloaded from the hg19 human genome assembly on the UCSC Genome browser. The scores used in Fig. 1E and Fig. S1B are derived from the total median expression level across all tissues (range: 0-1000). The GTEx data from the same source was also used to generate the tissue-specific expression plots Fig. S2). Genes were classified using the geneType field in the table, but some classes were merged as follows: all classes containing the word “pseudogene” were merged into our pseudogene set and classes “lincRNA”, “processed_transcript”, “antisense”, “macro_lncRNA”, “bidirectional_promoter_lncRNA” were merged into our lncRNA set.

The exon coordinates were also used to interrogate the GC percent content in 5-base windows track downloaded from the hg38 human genome assembly on the UCSC Genome browser (gc5base).

The rate of evolution of SNHGs was calculated by interrogating the 100-vertebrate conservation PhyloP track from the hg38 human genome assembly on the UCSC Genome browser with the exon coordinates for all SNHG lncRNA transcripts as well as coding gene ACTB as a reference. Since PhyloP scores are basewise -log10(p-values) of conservation (positive scores) or acceleration (negative scores)^21^ we used cutoffs of +2 for conserved positions or -2 for positions undergoing accelerated evolution; any nucleotide with a score between 2 and -2 was considered to be evolving neutrally. For each gene, we then calculated the average percentage of nucleotides under the three different evolutionary regimes across all transcripts.

### Primary human keratinocyte culture

Primary male human keratinocyte (strain Km) isolated from neonatal foreskin or night monkey (*Aotus trivirgatus*) keratinocytes isolated from an oesophageal biopsy^57^ were cultured at 37°C on mitotically inactivated 3T3-J2 cells in complete FAD medium (Gibco, 041-96624), containing one part Ham’s F12, three parts Dulbecco’s modified eagle medium (DMEM) (Gibco, 41966), 100 µM adenine, 10% (v/v) Foetal Bovine Serum (FBS), 1.8 mM CaCl_2_, 0.5 mg/ml hydrocortisone, 5 mg/ml insulin, 0.1 nM cholera toxin and 10 ng/ml Epidermal Growth Factor (EGF), as described previously^58^. Mouse keratinocytes were isolated from back skin and cultured in the same way except no CaCl_2_ was added to the medium (low Ca^2+^ FAD). Primary keratinocytes were used in experiments at passage 4-7. When subculturing or seeding cells for an experiment, the disaggregated keratinocytes were filtered through a nylon strainer to remove cell clumps and large differentiated cells.

Before mitotic inactivation, 3T3-J2 cells were cultured in DMEM (Gibco, 41966), supplemented with 10% bovine serum.

### siRNA transfection

Reverse transfection of siRNAs, miRNA inhibitors or biotinylated miRNAs was performed using INTERFERin transfection reagent (PolyPlus-transfection, 101000016) in accordance with the manufacturer’s instructions. All siRNAs were purchased from Horizon Discovery. siRNAs were diluted in OptiMEM medium (Gibco, 31985) and mixed with an appropriate volume of INTERFERin, depending on the transfection vessel. The siRNA/reagent complexes were allowed to form for 15 min at room temperature, before addition of the keratinocytes in Keratinocyte-Serum Free Medium (KSFM) (Gibco, 17005) supplemented with 0.15 ng/ml EGF and 30 mg/ml Bovine Pituitary Extract (BPE). The final concentration of the siRNA, unless otherwise specified, was 30 nM in the case of single transfections and up to a total concentration of 50 nM in the case of co-transfections of siRNA and miRNA inhibitors. For transfection of human or monkey keratinocytes tissue culture plastic vessels were coated with 20 µg/ml rat Collagen I (Sigma, C3867) overnight at 4°C. For transfection of mouse keratinocytes, tissue culture plastic vessels were coated with extracellular matrix deposed by feeder cells that were removed before keratinocyte addition. Four hours after transfection, the medium was changed to fresh complete KSFM in the case of human or monkey cells or to low Ca^2+^ FAD in the case of mouse cells. All knockdown experiments shown are at least three independent transfections.

### Clonogenicity assays

After siRNA transfection or plasmid stable lentiviral transduction, 500 (human), 1000 (night monkey) or 5000 (mouse) keratinocytes were plated on a 3T3 feeder layer per well of a six-well dish. After 12 days, feeders were removed, and keratinocyte colonies were fixed in 4% paraformaldehyde (Sigma, 158127) for 10 min then stained with 1% Rhodanile Blue (1:1 mixture of Rhodamine B (Sigma, R6626) and Nile Blue chloride (Sigma, 222550)). The number of colonies was counted manually. Unless otherwise specified, all statistics presented are calculated using well-level data. For siRNA transient transfections data was collected from at least 3 independent transfections, with the exception of Fig. 2E, which was generated from 2 independent transfections.

### RNA isolation, cDNA synthesis and quantitative Polymerase Chain Reaction (qPCR)

For mRNA quantification, RNA was extracted using the RNeasy Mini kit (Qiagen, 74104) and subsequently reverse transcribed using the QuantiTect Reverse Transcription (Qiagen, 205311) kit according to manufacturer’s instructions. When mRNA and snoRNA quantification from the same samples was required, RNA was extracted using the miRNeasy Mini kit (Qiagen, 217004) and subsequently reverse transcribed using the mScript II RT kit (Qiagen, 218161) with HiFlex Buffer according to the manufacturer’s instructions. The cDNA thus obtained was diluted to 2.5-5 ng/µl and specific targets were amplified by qPCR using the Fast SYBR® Green Master Mix (Applied biosystems, 4385612). Expression of all targets was normalised against the expression of three reference genes (RPL13A, ATP5B and TBP) (ΔCq). For miRNA quantification, RNA was extracted using the miRNeasy Mini kit (Qiagen, 217004) and subsequently reverse transcribed using the miRCURY LNA RT kit (Qiagen, 339340) according to manufacturer’s instructions. The cDNA thus obtained was diluted to 2 ng/µl and specific target miRNAs were amplified by qPCR using the miRCURY LNA SYBR Green PCR kit (Qiagen, 339346). Expression of all targets was normalised against the expression of two reference genes (SNORD48 and U6) (ΔCq). Where indicated, expression was also normalised against control samples (ΔΔCq).

### smRNA (F)ISH and analysis of tissue sections and cultured cells

Chromogenic *in-situ* hybridisation of lncRNA on cultured cells was performed using the RNAscope 2.5 HD Assay – RED kit (ACD, 322350) according to the manufacturer’s instructions. Fluorescent *in-situ* hybridisation of lncRNA and mRNA on tissue sections was performed using the RNAscope Multiplex Fluorescent v2 kit (ACD, 323100) according to the manufacturer’s instructions. Chromogenic *in-situ* hybridisation of miRNA on cultured cells was performed using the miRNAscope HD Reagent Kit - RED (ACD, 324500) according to the manufacturer’s instructions.

For the quantifications, in the case of cultured cells, the number of individual dots in at least 180 cells/staining was counted. In the case of tissue sections, fluorescence intensity was scored by using the nucleus detection algorithm on QPath 0.3.2 and expanding the area around the nucleus by 2 µm. Epidermal layers in Fig. 3D were annotated manually. Positive cell positioning shown in Fig. 3F was quantified by manually thresholding SNHG7 intensity and subsequently scoring the Y position of the positive cells relative to the local minimum and maximum Y coordinates of the epidermis. The local minima and maxima, used in order to account for ondulations in the epidermis were calculated as the highest and lowest cell position every 20 consecutive cells on the X axis.

### Wound/re-epithelialisation assay of skin biopsies

Surplus surgical waste skin was obtained from consenting patients undergoing plastic surgery. This work was ethically approved by the National Research Ethics Service (UK) (HTA Licence No: 12121, REC No: 14/NS/1073). The tissue was sterilised, washed several times with PBS and cut into 1 cm^2^ pieces. Partial thickness wounds, comprising the epidermis and upper part of the dermis, were created with a 4 mm punch biopsy. Skin explants were then placed into 6-well hanging cell culture inserts (Millipore) and FAD medium added to create an air–liquid interface. *Ex vivo* explants were maintained in culture with FAD medium and an air–liquid interface for 2 weeks with media changes every 48 h.

### Immunocytofluorescence

Plates were washed once with PBS and fixed with 4% Paraformaldehyde (Sigma, 158127) incubated for 10 min at room temperature. Plates were then permeabilised by incubating with 0.2% Triton-X-100 in PBS for 5 min at room temperature, incubated with blocking buffer (10% FBS, 0.25% Fish Skin Gelatin in PBS) for 1 h and stained overnight at +4°C with the primary antibodies anti-Involucrin (SY3 or SY7 clones), anti-cleaved Caspase 3 (Asp175) (Cell Signalling 9661), anti-Ki67 (abcam, ab16667), anti-Integrin β1 (eBioscience, 14-0299-82), anti-Keratin 10 (BioLegend, 905404), anti-Keratin 14 (Biolegend, 906004) diluted to 0.5-1 µg/ml or according to manufacturer’s instructions in blocking buffer. Plates were then washed three times with PBS, stained with the secondary antibodies AlexaFluor 555 donkey anti-mouse (Invitrogen, A32773), AlexaFluor 647 goat anti-chicken IgY (Invitrogen, A21449) or AlexaFluor 488 donkey anti-rabbit (Invitrogen, A21206) at 1 µg/ml, and the nuclear dye DRAQ5 (abcam, ab108410) at 10 µM or DAPI at 0.1 µg/ml in blocking buffer. Secondary stains were incubated for 2 h at room temperature protected from light and plates were finally washed three times with PBS before being imaged using the Perkin-Elmer Operetta High-Content Imaging System.

### High content imaging analysis

Images acquired with the Perkin-Elmer Operetta High-Content Imaging System were analysed using custom algorithms in the Perkin-Elmer Harmony high-content analysis software package. Nuclei were initially defined using the DRAQ5 channel; small (< 2000 µm^2^) and highly irregular (roundness < 0.6) nuclei were excluded from the analysis; to minimise the misattribution of the cytoplasm areas, the level of cytoplasmic staining was inferred from the fluorescence intensity in a ring around the nucleus, as Involucrin staining was homogeneous throughout the cytoplasm. Terminally differentiating cells were identified by manual thresholding of the of Involucrin perinuclear fluorescence intensity. The complete Harmony image analysis sequence is available on request.

### OP-Puro assay

General protein synthesis levels were measured using the Click-iT Plus OPP Alexa Fluor 488 Protein Synthesis Assay Kit (Invitrogen, C10456). After 48 h of siRNA transfection, cells were treated with cycloheximide or control diluent for 2 h before O-propargyl-puromycin treatment for 30 min, fixation in 4% paraformaldehyde and staining according to the manufacturer’s instructions.

### Skin reconstitution on de-epidermised dermis (DED)

Dermis from excess human adult surgical waste skin was decellularized by repeat freeze/thaw cycles and used as a substrate for new epidermis formation. Keratinocytes transfected with siRNA targeting SNHG7 or a scrambled control were transferred on decellularized dermal substrates 24 h after transfection and kept growing on a tissue culture insert in contact with FAD medium conditioned by a feeder layer at the air-liquid interface for 2 weeks. Dermal substrates were subsequently embedded in optimal cutting temperature compound (OCT, Life Technologies) and frozen before sectioning and staining with haematoxylin and eosin. Epidermal thickness was scored across the entire length of multiple sections per DED and the mean thicknesses of two DEDs seeded with two independent transfections per condition were compared.

### Lentiviral transduction of human keratinocytes

Overexpression of wild-type or mutant human SNHG7 in primary human, night monkey or mouse keratinocytes was achieved by lentiviral transduction. Lentiviral particles containing control (“GFP only”), wt human SNHG7 (transcript NR_024542, “hSNHG7”) or mutant human SNHG7 lncRNAs (“hSNHG7mut”, see Fig. 6A and Fig. S9A) custom overexpression plasmids (Oxgene) were transduced onto keratinocytes for 24 h at a multiplicity of infection of 3 m.o.i. in the presence of 5 µg/ml polybrene. Both control and overexpression plasmids encoded GFP as an indicator of transduction efficiency. Stably transduced cells were FACS-sorted 48h after transduction and the GFP-positive cells were subcultured and expanded. GFP transcription was under the control of a separate promoter to avoid the generation of a hybrid RNA sequence that may have interfered with SNHG7 function.

### Suspension-induced keratinocyte differentiation

Keratinocytes were differentiated in suspension as described^27,59^. Pre-confluent cultures were disaggregated in trypsin/EDTA and resuspended at a concentration of 10^5^ cells/ml in medium containing 1.45% methylcellulose. Aliquots were plated in 6-well plates coated with 0.4% polyHEMA; this ensured that there was no cell-substratum adhesion. The suspended cells were subsequently incubated at 37°C. At each collection time point, the methylcellulose was diluted with PBS and the cells recovered by centrifugation.

### 3’ and 5’ RACE

Sequencing of *Aotus trivirgatus* SNHG7 lncRNA was achieved by amplifying the 3’ and 5’ end of the cDNA from internal primers using the 5’/3’ RACE kit, 2^nd^ Generation (Roche, 03353621001). Since no genome data was available for *Aotus trivirgatus*, internal primers were designed from the sequence of the closely related species *Aotus nancymaae*.

### Sequence alignment

*A. trivirgatus* SNHG7 lncRNA sequences were aligned to the human transcript (NR_024542) by using a supervised application of the Needleman-Wunsch global alignment algorithm on an exon-by-exon basis. The intron-exon structure of the *A. trivirgatus* gene was inferred by manually comparing the genomic sequences of *H. sapiens* and *A. nancymaae* (NW_018496780) for which an assembled genome was available.

The *M. musculus* SNHG7 sequence (NR_024068) was aligned to the human transcript using a supervised application of the Needleman-Wunsch global alignment algorithm on an exon-by-exon basis using the annotated exons.

Alignments were visualised using the Geneious Prime software (Geneious).

### RNAseq analysis

Samples transfected with siRNA targeting SNHG7 or a scrambled control were harvested 24 h and 48 h post-transfection and sent to Azenta Life Sciences for bulk RNA sequencing. Sequence reads were trimmed to remove possible adapter sequences and nucleotides with poor quality using Trimmomatic v.0.36. The trimmed reads were mapped to the Homo sapiens GRCh38 reference genome available on ENSEMBL using the STAR aligner v.2.5.2b. Unique gene hit counts were calculated by using featureCounts from the Subread package v.1.5.2. Only unique reads that fell within exon regions were counted.

After extraction of gene hit counts, the gene hit counts table was used for downstream differential expression analysis. Using DESeq2, a comparison of gene expression between the knockdown (siSNHG7) and control (siScramble) samples was performed at 24 h and 48h post transfection. The Wald test was used to generate p-values and log_2_ fold changes. The p-values were then adjusted for multiple testing using the Benjamini and Hochberg method. The adjusted p-values for this test are referred to in the manuscript as p_adj_.

GO term enrichment analysis was performed using ShinyGO 0.77^60^, inputting all significantly differentially expressed genes (p_adj_ < 0.05) at each time point and assessing enrichment of Biological Process GO terms with respect to all detected genes in the RNA-seq.

Significantly (p_adj_ < 0.05) downregulated genes at each time point were scanned for enrichment of miRNA response elements (MREs) against the miRTarBase database (version 7.0) of experimentally validated miRNA-mRNA interactions using MIENTURNET^35^. Briefly, this web tool performs a hypergeometric test to assess the probability of finding a certain number of interactions (or more) between a miRNA and a list of target genes, considering the total number of interactions that miRNA engages in, and the total number of miRNA-target interactions present in the database, according to the formula:

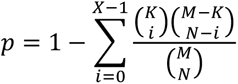

Where M is the universe of total interactions encompassed in the miRTarBase database, K is the total number of interactions miRNA i engages in, N is the number of genes in the candidate list (in our case the significantly downregulated genes following SNHG7 knockdown), and X is the number of miRNA interactions found within the candidate list. The p-values thus calculated are subsequently adjusted for multiple testing using the Benjamini-Hochberg method. The adjusted p-values for this test are referred to as FDR in the manuscript.

The same tool was also used to assess enrichment of REACTOME Pathways for the downregulated targets of each candidate SNHG7-interacting miRNA.

### Intracellular fractionation

Isolation of cytoplasmic and nuclear fractions from human keratinocytes was performed following a published protocol^61^. After fractionation and RNA extraction, all samples were diluted to the same RNA concentration and 250 ng of RNA from each sample spiked with 100 ng mouse RNA to control reaction efficiency were reverse transcribed using the miScript II RT kit (Qiagen, 218161) with HiFlex Buffer according to manufacturer’s instructions. Amounts in each fraction were then calculated by multiplying the RNA found in 250 ng by the total volume of each fraction.

### RNA pulldown

Pulldown of biotinylated microRNAs was performed as described in the literature^62^ 24 h after transfection with biotinylated miRNAs purchased from Integrated DNA technologies. Before pulldown, 1/10^th^ of each sample was aliquoted to serve as the input reference and the RNA pulled down in each sample was normalised to the corresponding input.

### MRE conservation analysis

In order to assess the presence and conservation of miRNA response elements (MREs) in SNHGs we used the “maximum exonic coverage” of all transcripts that did not overlap coding sequences or snoRNAs for each of the SNHGs.

First, SNHG transcripts that did not contain exons overlapping coding genes or snoRNAs were selected and the coordinates for their exons were extracted from the hg19 human genome assembly on the UCSC Genome browser. Second, the exonic coordinates were used to extract the sequences from the 100-vertebrates multiple sequence alignment (multiz100way) from the hg19 human genome assembly on the UCSC Genome browser. Third, the extracted sequences were stitched together using the minimum and maximum coordinate of all selected transcripts for each SNHG. This generated, for every SNHG, a multiple sequence alignment of all exonic portions of the gene stitched together in an idealised “maximum exonic coverage” transcript. Fourth, we used the TargetScan 7 suite to identify MREs in the species present in the multiple sequence alignment and to calculate their branch length scores. For this analysis, we did not require miRNAs to have been annotated in each species but only seed sequences of deeply conserved miRNAs were used. Since not all individual isoforms are analysed, it is possible that some additional miRNA sites can be created at alternatively spliced junctions.

In order to compare the distributions of MRE conservation between SNHGs and the 3’UTRs of coding genes, we performed the same MRE conservation analysis on 250 UTRs maintaining the same background conservation binning^44^ distribution we had for the SNHG set.

Some of the steps in this analysis were performed using the Galaxy platform^63^.

## Author contributions

This study was conceived by MVR and FMW. MVR designed and performed all experiments and analyses unless otherwise stated. KHS contributed to experimental design, isolated the mouse keratinocytes, and imaged the psoriasis and wound sections. CP performed the biopsy wound/re-epithelialisation assay. CG imaged the smRNA FISH skin sections and helped with data analysis. VAN assisted with scRNAseq data processing and analysis. The manuscript was written by MVR and edited by KHS and FMW with contributions from all co-authors.

## Acknowledgements

MVR would like to thank Dr Ajay Mishra for early insights about this project and Dr Flavia Michelini for useful advice about biotin-mediated RNA pulldown experiments. FMW gratefully acknowledges financial support from Cancer Research UK (C219/A23522), the Medical Research Council (G1100073) and the Wellcome Trust (096540/Z/11/Z; 211276/E/18/Z). We are also grateful to the National Institute for Health Research (NIHR) Biomedical Research Centre based at Guy’s and St Thomas’ NHS Foundation Trust and King’s College London. The views expressed are those of the author(s) and not necessarily those of the NHS, the NIHR or the Department of Health.

---

**Fig. S1.**
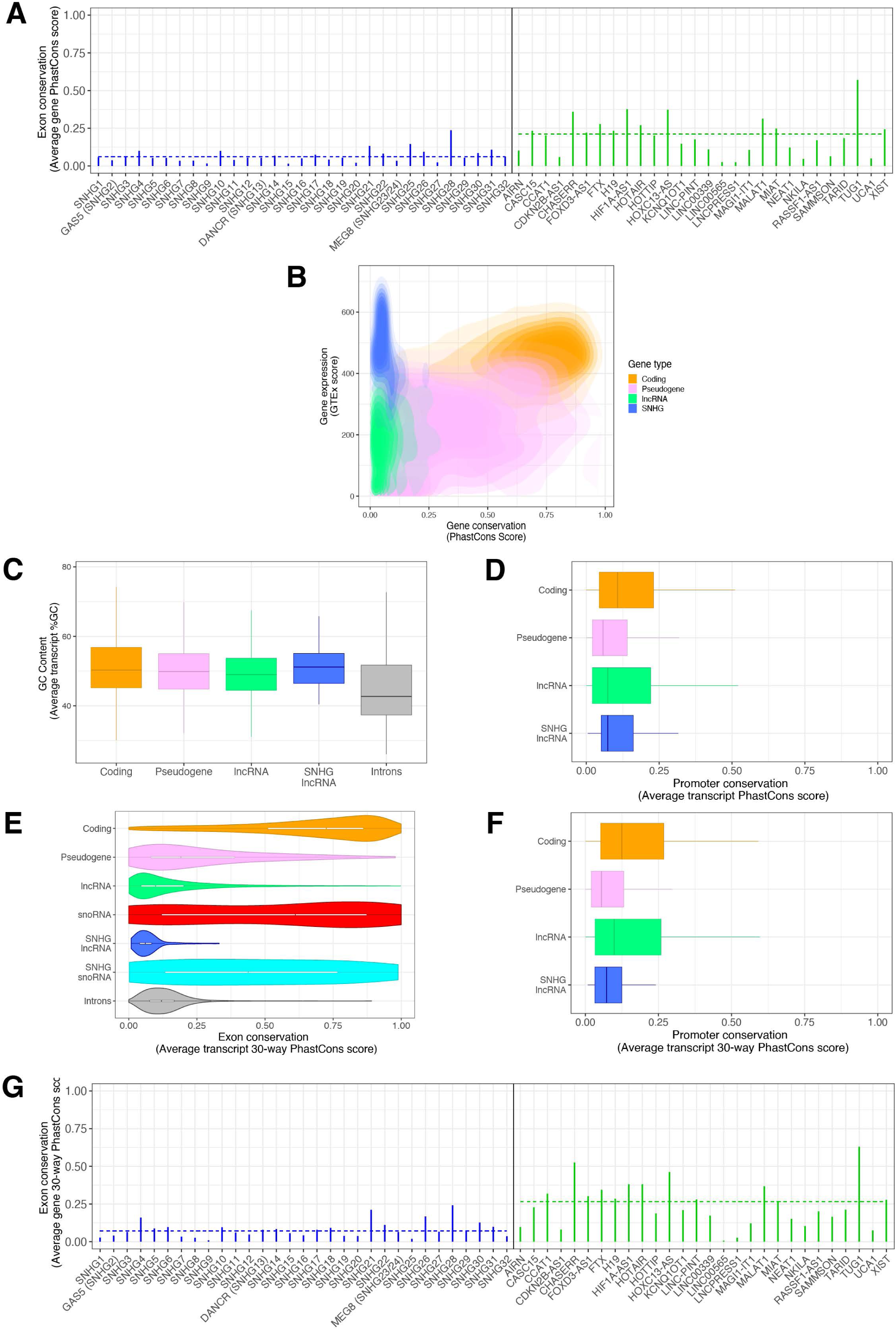
SNHGs in the genomic landscape. (**A**) Exonic conservation of SNHGs (blue) compared to a set of lncRNA with characterised functions (green). Dashed lines represent the mean conservation of each group. (**B**) Conservation and expression distribution of different transcript classes genome wide. (**C**) Distribution of GC content in SNHG lncRNA compared to other classes of transcripts genome-wide. (**D**) Conservation of promoters in SNHGs and other gene classes. (**E**) Conservation of different classes of transcripts genome-wide evaluated by using PhastCons scores generated from alignment of 30 mammalian genomes (28 primates). (**F**) Conservation of promoters in SNHGs and other gene classes evaluated by using PhastCons scores generated from alignment of 30 mammalian genomes (28 primates). (**G**) Exonic conservation of SNHGs (blue) compared to a set of lncRNA with characterised functions (green) evaluated by using PhastCons scores generated from alignment of 30 mammalian genomes (28 primates). Dashed lines represent the mean conservation of each group. All boxplots indicate the median and the interquartile range.

**Fig. S2.**
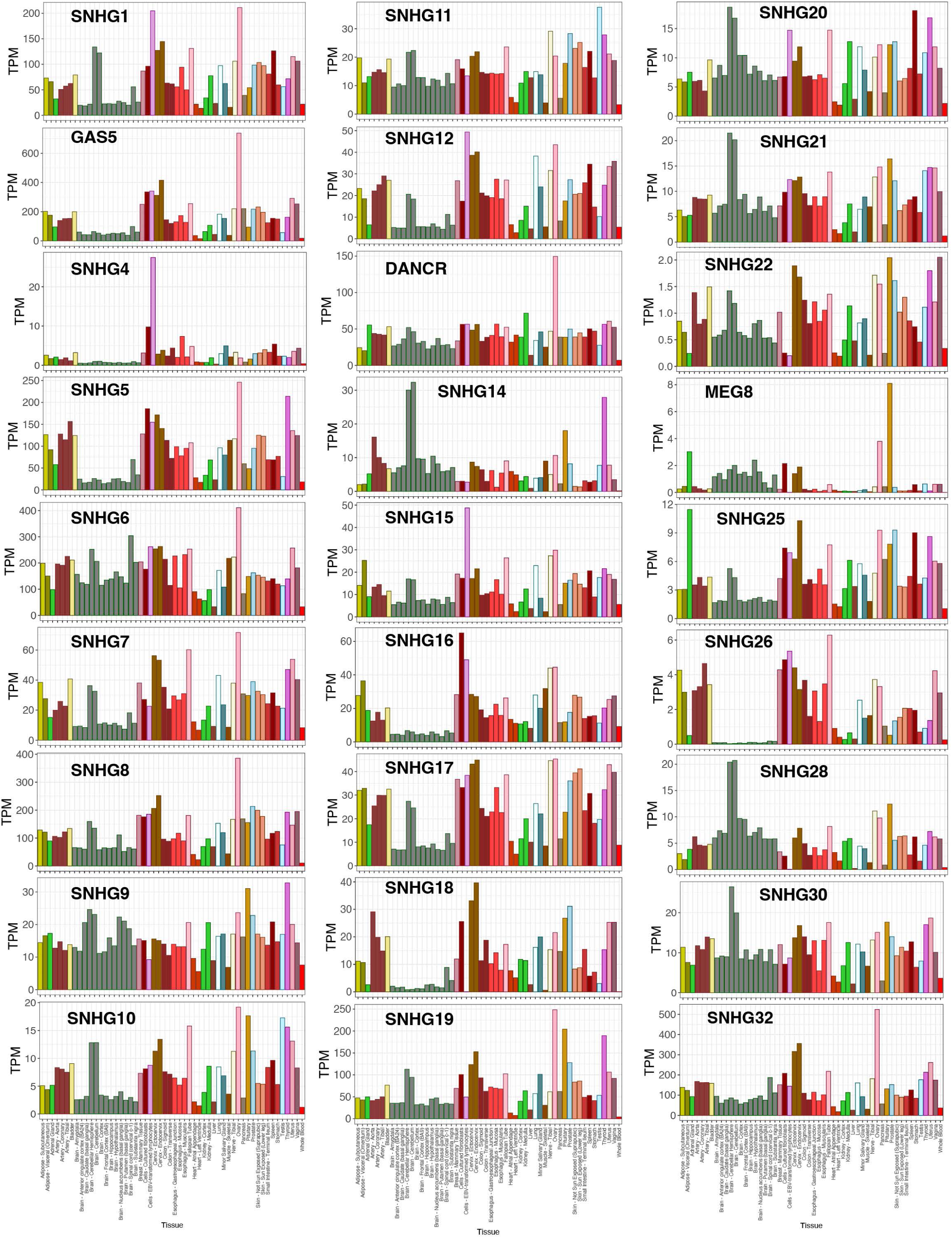
**SNHGs expression across human tissues.**

**Fig. S3.**
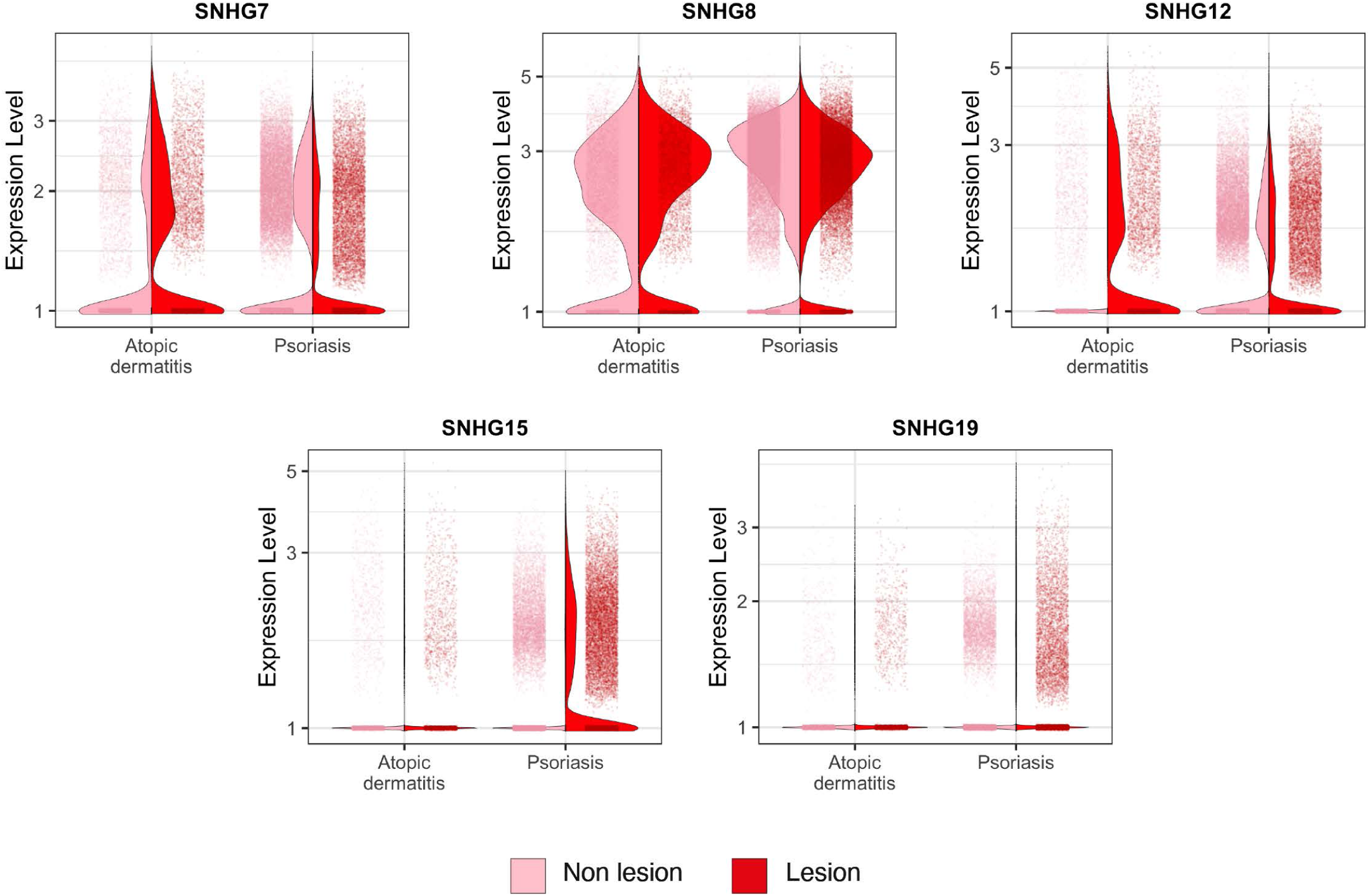
Poorly conserved SNHGs are regulated in skin inflammatory conditions. Expression of five epidermal SNHGs in scRNA-seq data from the lesional and non-lesional areas of atopic dermatitis, and psoriasis-affected tissues.

**Fig. S4.**
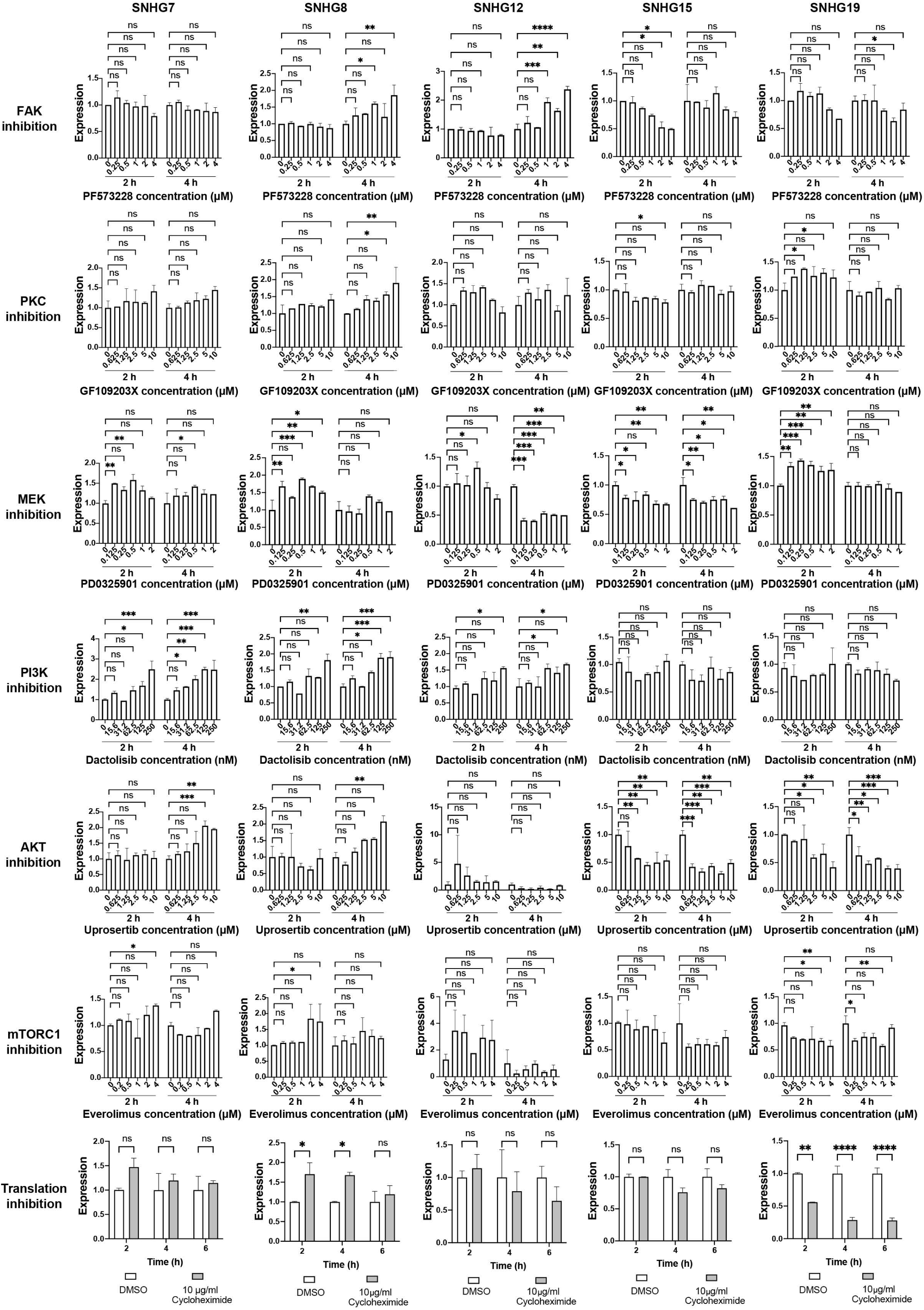
Epidermal SNHGs expression is controlled by multiple signalling pathways. Changes in the expression of five epidermally expressed SNHGs in primary human keratinocytes in response to treatment with increasing doses of different pathway inhibitors for 2 h or 4 h. Data shown are mean +/- SD. Dunnet’s multiple comparisons test, n ≥ 2. **** p < 0.0001, *** p < 0.001, ** p < 0.01, * p < 0.05.

**Fig. S5.**
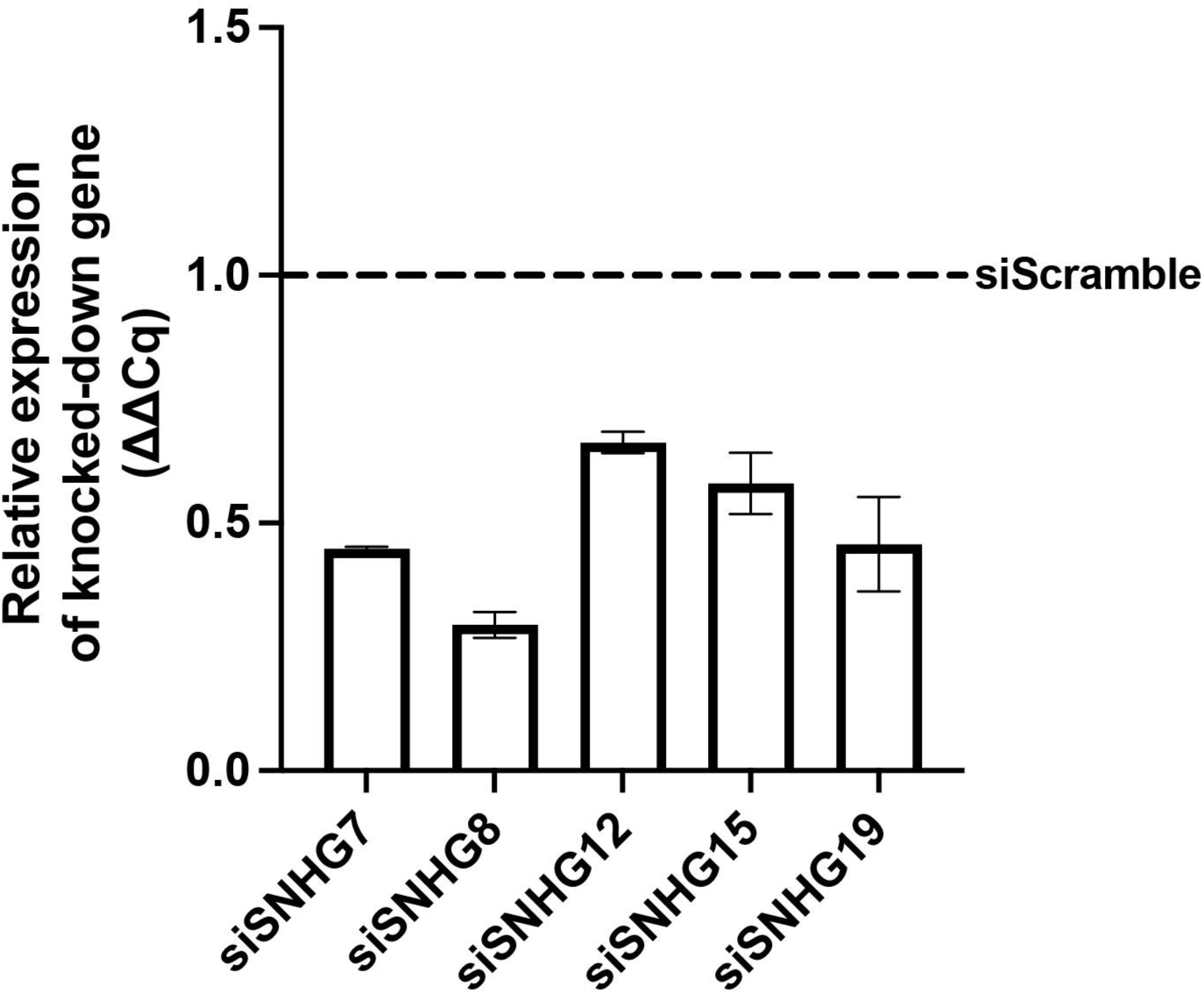
**SNHG knockdown efficiency 24 h post-transfection in clonogenicity assays.**

**Fig. S6.**
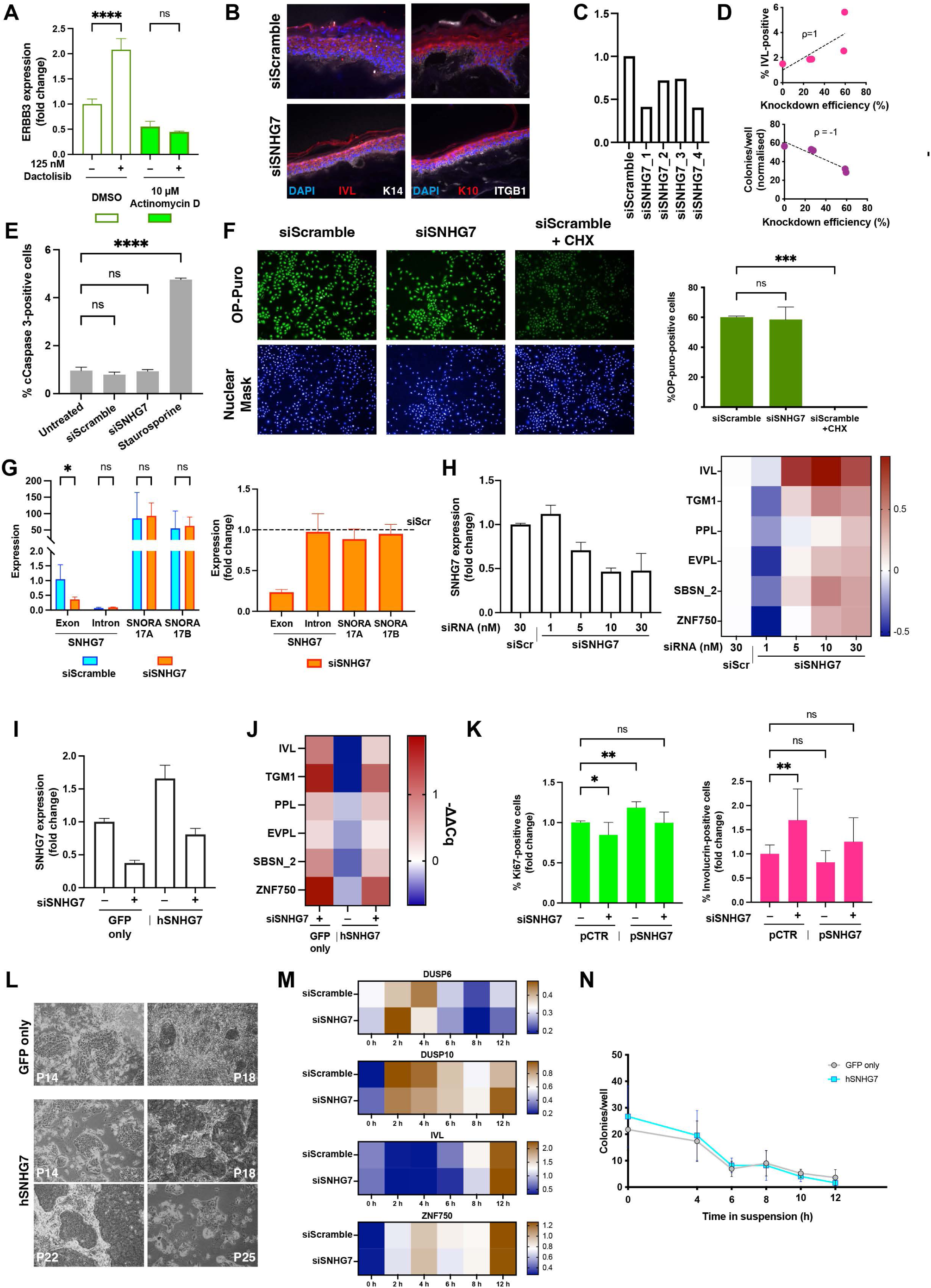
SNHG7 lncRNA affects epidermal stem cell proliferation and differentiation. (**A**) ERBB3 expression after 4 h PI3K inhibition with or without prior 1 h treatment with a transcription inhibitor. n = 5 independent treatments. (**B**) Epidermal differentiation marker staining of epidermis reconstituted by control or SNHG7 knockdown cells. (**C**-**D**) Deconvolution of the siRNA pool targeting SNHG7. Transfection efficiency of the individual siRNAs (**C**) and correlation between knockdown level and Involucrin expression (**D**, top) or clonogenic capacity (**D**, bottom). (**E**) Effect of SNHG7 knockdown on apoptosis. Quantification of the percentage of cells staining positive for cleaved Caspase 3. n = 2 independent transfections/treatments. (**F**) Effect of SNHG7 knockdown on translation. Representative images (left panels) and quantification (right bar plot) of keratinocytes stained with OP-Puro in control conditions, after SNHG7 knockdown and after treatment with translation inhibitor cycloheximide (CHX). n = 2 independent treatments/transfections. (**G**) Effect of siSNHG7 treatment on different transcripts arising from the locus. Shown are expression levels relative to reference genes (left) and change in expression after knockdown relative to control (right). n = 2 independent transfections. (**H**) Titration of siSNHG7 to identify the minimal effective concentration. Knockdown efficiency at 48 h post transfection (left) and differentiation marker expression heatmap (right). (**I**-**K**) Rescue of the SNHG7 knockdown phenotype by lncRNA overexpression. Shown are SNHG7 lncRNA expression levels in all samples (**I**), differentiation marker expression 72 h post-transfection relative to control (**J**), and quantification of differentiation and proliferation marker staining 96 h post-transfection (**K**). n = 8 independent transfections. (**L**) Representative images of cultured keratinocytes overexpressing SNHG7 or GFP only at different passages. Cells were transduced at the same time and were subcultured seeding the same number of cells at each passage. (**M**) Time course of commitment (DUSP6, DUSP10) and differentiation (ZNF750, IVL) marker expression after suspension-induced differentiation of control and SNHG7 knockdown cells. pCTR, control plasmid; pSNHG7, overexpression plasmid. (**N**) Suspension-induced differentiation time course of control and SNG7-overexpressing keratinocytes. Scale bars, 50 µm. Data shown in all bar plots are mean +/- SD. (**A**) Šidák’s multiple comparisons test. (**E**) (**F**) and (**K**) Dunnet’s multiple comparisons test. (**G**) Multiple unpaired two-tailed t-tests.

**Fig. S7.**
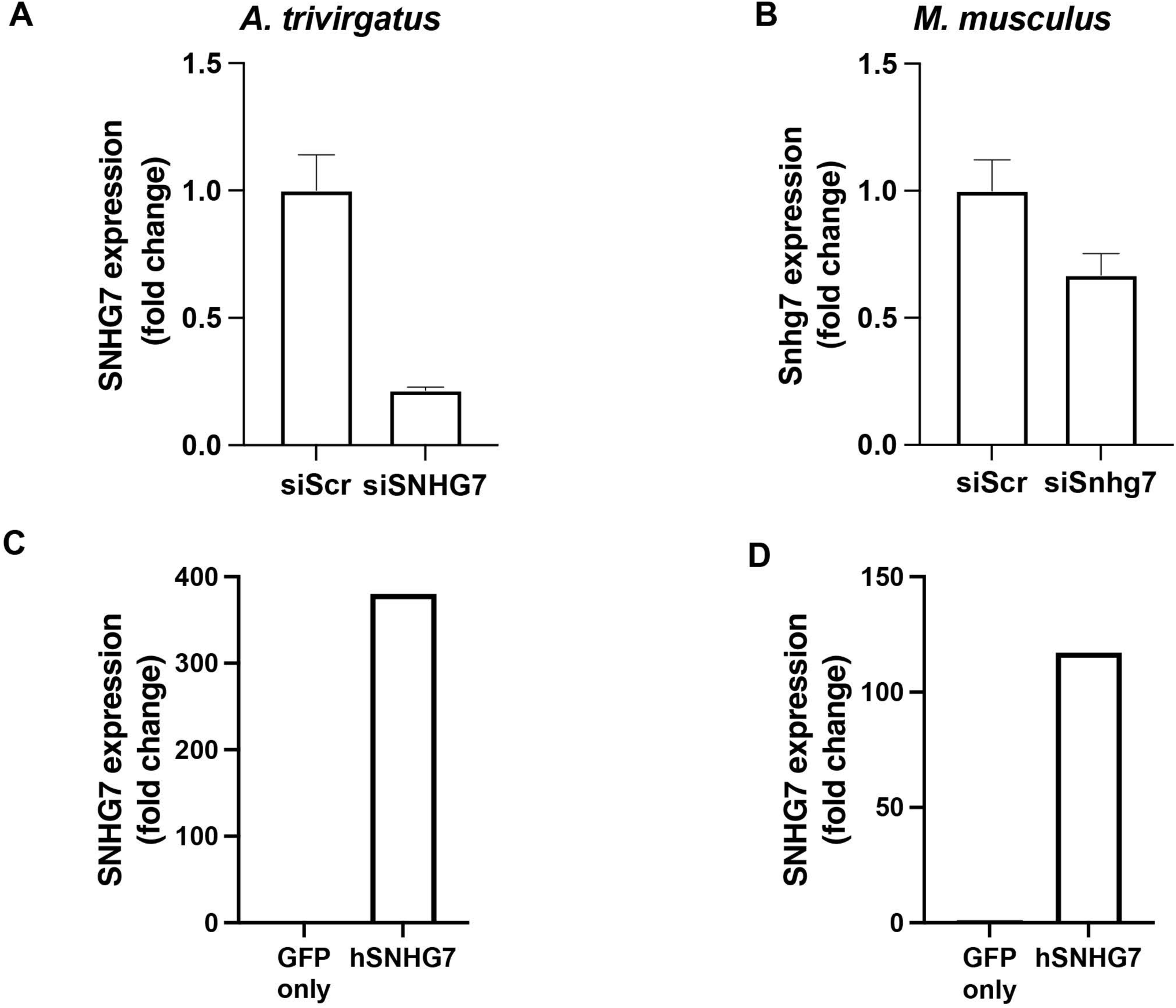
SNHG7 knockdown and overexpression efficiency in A. trivirgatus and M. musculus cells. (**A**-**B**) Expression of endogenous SNHG7 after 48h siRNA transfection in A. trivirgatus (**A**) or M. musculus (**B**) keratinocytes. (**C**-**D**) Expression of human SNHG7 in *A. trivirgatus* (**C**) or *M. musculus* (**D**) keratinocytes after stable transfection with an overexpression plasmid.

**Fig. S8.**
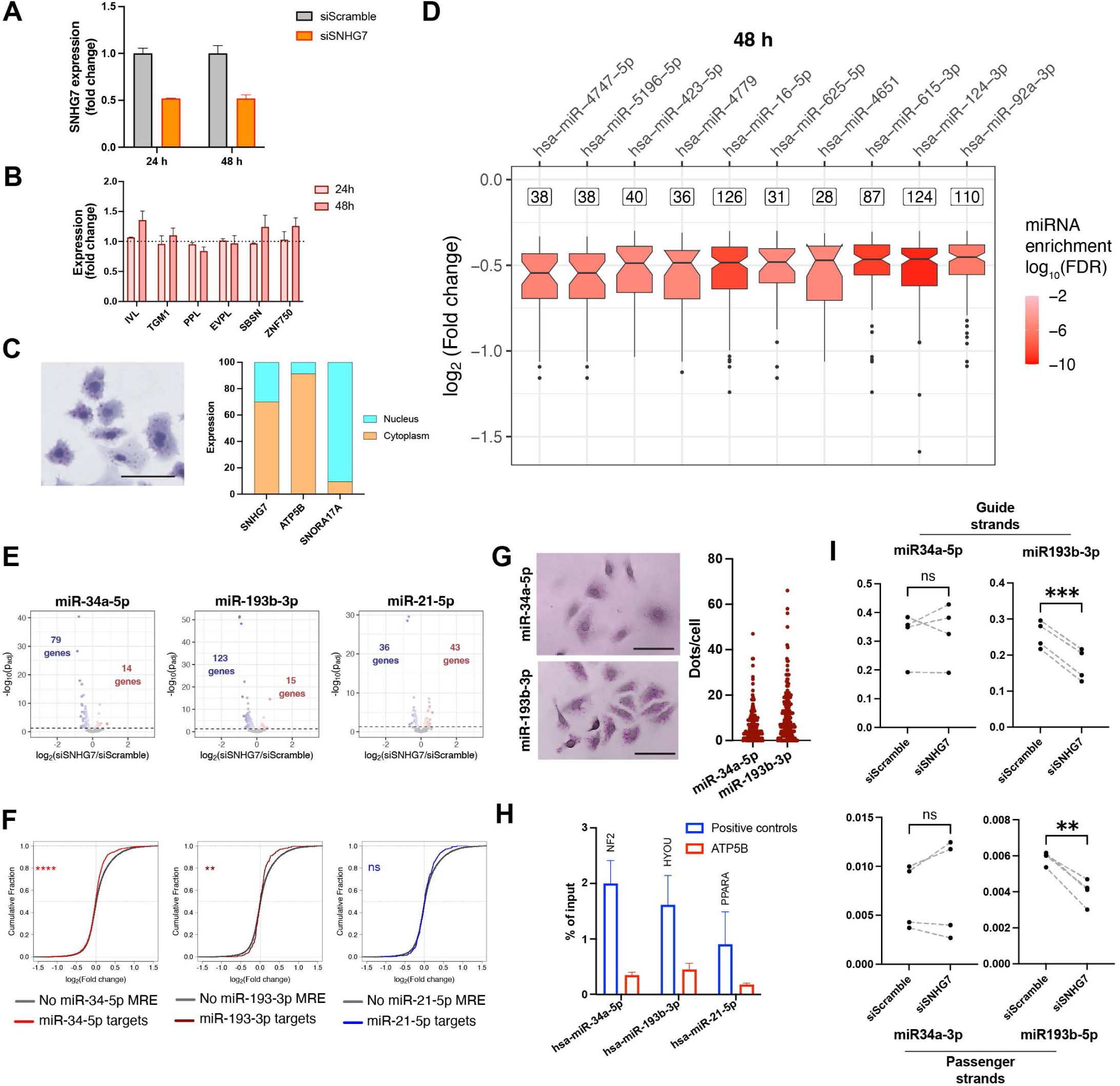
SNHG7 lncRNA affects the expression of targets of interacting miRNAs. (**A**) Knockdown efficiency in the RNA-seq. (**B**) Expression variation of selected differentiation markers in the RNA-seq. (**C**) Intracellular distribution of SNHG7 assessed by smRNA cytochemistry (left) or subcellular fractionation (right). Subcellular distributions of a mRNA (ATP5B) and a snoRNA (SNORA17A) are used as controls and references. (**D**) miRNA response element enrichment in significantly (padj < 0.05) downregulated genes 48 h post-transfection. The number of target genes for each miRNA is shown above the box plot of their differential expression. miRNAs are sorted based on the median downregulation of their target genes. The colour of the boxes indicates the significance of the enrichment. (**E**) Validated miRNA target genes in the siSNHG7 RNAseq data. Volcano plots show all validated target genes for the two candidate miRNAs (miR-34a-5p and miR-193b-3p), and a control miRNA (miR-21-5p). (**F**) Cumulative distributions of the gene expression change between control and knockdown cells for predicted targets of miR-34a (red, left), miR-193b (darkred, centre) or miR-21 (blue, right) compared to transcripts that do not contain the respective MREs (grey). Kolmogorov-Smirnov test. (**G**) smRNA cytochemistry and quantification of the expression of two candidate miRNAs in primary human keratinocytes. Lines indicate the median. (**H**) Enrichment of control genes after pulldown of biotinylated miRNAs. Genes containing MREs for the indicated miRNAs are used as positive controls for the pulldown and are indicated above the bars. ATP5B was used as a negative control as it does not contain MREs for any of the miRNAs. (**H**) Variation in the levels of guide and passenger strands of candidate miRNAs after SNHG7 knockdown. Paired two-tailed t-tests. Pairs of points (connected by dotted grey lines) represent individual experiments. Data shown in all bar plots are mean +/- SD. **** p > 0.0001 *** p > 0.001, ** p > 0.01.

**Fig. S9.**
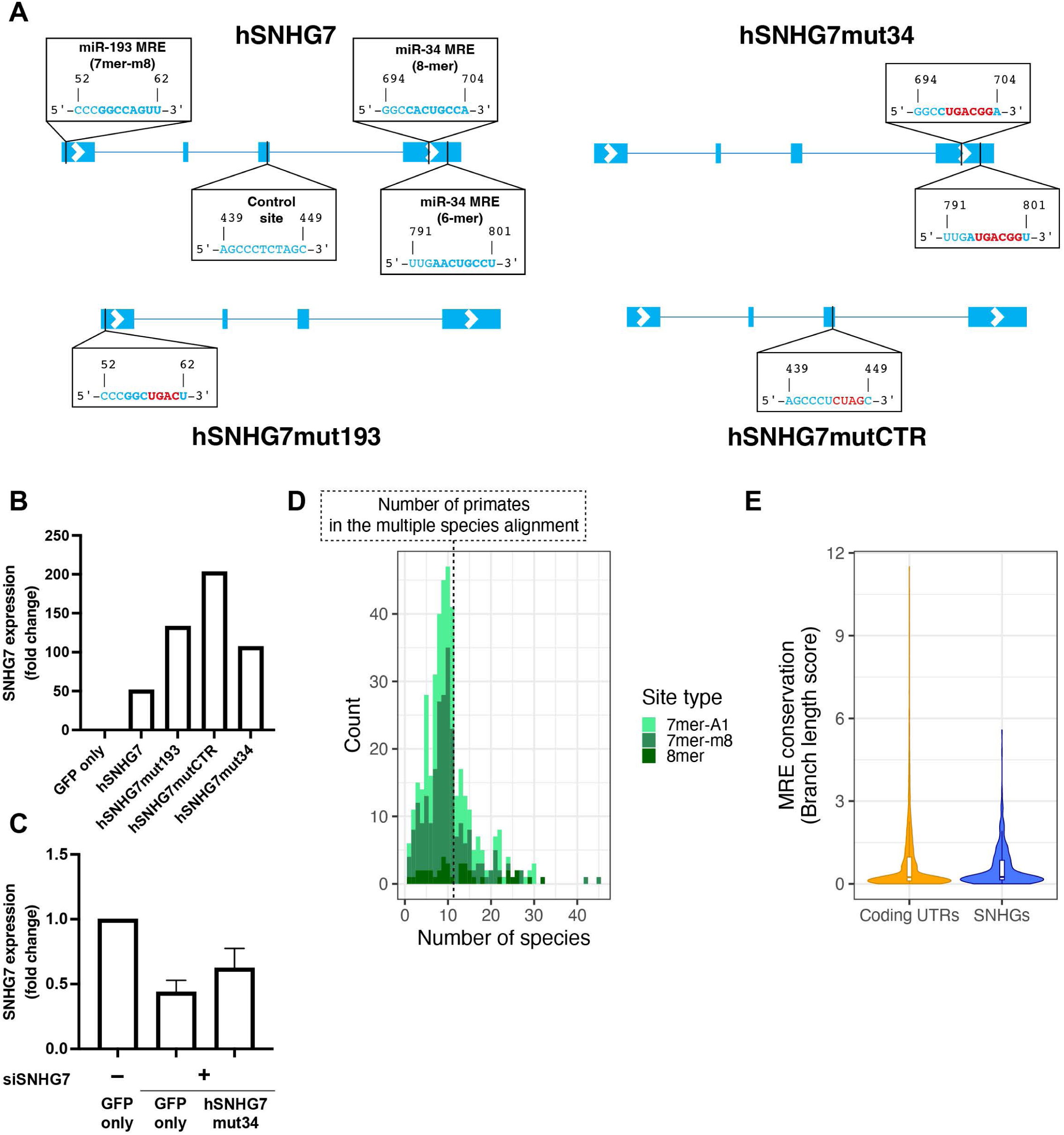
Evolution of SNHG-MRE interactions. (**A**) Detailed schematic of the mutant transcripts used in overexpression/rescue experiments. (**B**) Expression of human SNHG7 mutants in *A. trivirgatus* keratinocytes after knockdown stable transduction with an overexpression plasmid. (**C**) Expression of SNHG7 in human keratinocytes after knockdown in keratinocytes stably transduced with a control (GFP only) or SNHG7 mutant overexpression plasmid. (**D**) Distribution of the MREs present on human SNHGs according to the number of species that share them. MREs are broken down into the three different types of seed binding. A dashed line marks the number of primate species in the multiple alignment used for this analysis, indicating that all MREs shared by a larger number of species are conserved beyond the primate lineage. MREs shared by a smaller number of species can either be conserved only among primates or be conserved beyond the primate lineage but lost in one or more primate species. (**E**) Conservation of MREs for deeply conserved miRNAs found in SNHGs and in a set of 250 3’UTRs of coding genes.

## Notes

### Competing Interest Statement

The authors have declared no competing interest.

### Summary of Updates

This is a preliminary revision that addresses comments raised by the reviewers.

