## Supplementary Data 1 for "Neutral evolution of snoRNA Host Gene long non-coding RNA affects cell fate control"

>Aotus trivirgatus SNHG7 (transcript 1)

CTCTGCGCGCGCCGGCGGCTGCCATGGCGGGACGTCTGCTCACTGGAAACACACGTGCGAGGGGTCCTGGGGTGCTGGGAAGGAGGTGACTTAGCCTGTGATGGACTTCCAGTGTGAGCAGTGGCCAGAGTGACGAGGCCAACCGGCCCCAGTCCGAATCAAGATGCAGAGGCCAGGATGTGGGCGCAGCCCCGTGCCAAGAGGCGGGCTGGCATAGGACCTCCGGCACCCAGGCTGTTTGCGGCCTCAGAGCCCAGCTTTCCGCACGCCCACCTGCCCCCAGGGCCACGGTTGCAGCTCCTGCTCTGCCTGCATTCCAGGGATGGGCAGGCTGGCATCGGGACGCCCACCGCCTCTGCCTGGGTAGTGCTGTGTGTTCCAGCCGGCCAGGGCAGCTGCCAGGACCACCCCTCCATTTGAGTATCCCGGTTCTTAAGTTCTGCCATTGTGGTGTTCTGCTGGAAAAAGAACCATTTGGCTGTGTCTGAACTGCCTGGAACCCAAGATCCCGAGTTATTTTTTACTGTATTTGAGTCATCTTGTGTTTGTTGTTTTTACCCCAAGGGGAAAATCTAGATGGAAAACATTTATTTTAAAATACAGGATGAAGGGAATTAAAAGATTTAATGCACATTTCTTCAAGGATAGTATTTCTGTATTGGCAAAATTTGAGAATAAATGGGTCTGGAACGAAAAAAAAAAAAAAA

>Aotus trivirgatus SNHG7 (transcript 2)

CGCGTGAGCCGCGGGATGGGGGCCCGGGCCCGGGAGGAGGCGCCGTGCTGTGTCCCTCCCCGCGCGGTTTCCAGCCGGGAAGCTTCGGGAAGCCTGGTGAGGGCCGAGTGCGCGAATTCGGACCTAAGCGGAAAGCGCCCCGYCCACCCGCGCCCCTTCCGCCCCTGCTGCTGCGTCCCCGAGTCGCGGAGGCTCTGGGGACGTCGCCTCCTGTGTCGGCATCTTCGAGAAATGGATTTCTCGTGCCGTGTCCACGCGTCGGGTGTTTCCGTGTGACTGGCCGCTCAGCGGGAGGCTGTCCTGGCGGAAGGAGTCCGGTGACCCCCGGACTAAATACTGTTACAGGACAGGTGCGCGCCTGTCCTTGGGGGGCATCCGCCTCGTGGTCCTGGTCCCTGGACACCTGCAGGGGATCAGCCTTCCGTGGCCACTGTTGTGACTCAACTTCCCCTCCTGCAGATTAAGGAGAGAGACGTCTGCTCACTGGAAACACACGTGCGAGGGGTCCTGGGGTGCTGGGAAGGAGGTGACTTAGCCTGTGATGGACTTCCAGTGTGAGCAGTGGCCAGAGTGACGAGGCCAACCGGCCCCAGTCCGAATCAAGATGCAGAGGCCAGGATGTGGGCGCAGCCCCGTGCCAAGAGGCGGGCTGGCATAGGACCTCCGGCACCCAGGCTGTTTGCGGCCTCAGAGCCCAGCTTTCCGCACGCCCACCTGCCCCCAGGGCCACGGTTGCAGCTCCTGCTCTGCCTGCATTCCAGGGATGGGCAGGCTGGCATCGGGACGCCCACCGCCTCTGCCTGGGTAGTGCTGTGTGTTCCAGCCGGCCAGGGCAGCTGCCAGGACCACCCCTCCATTTGAGTATCCCGGTTCTTAAGTTCTGCCATTGTGGTGTTCTGCTGGAAAAAGAACCATTTGGCTGTGTCTGAACTGCCTGGAACCCAAGATCCCGAGTTATTTTTTACTGTATTTGAGTCATCTTGTGTTTGTTGTTTTTACCCCAAGGGGAAAATCTAGATGGAAAACATTTATTTTAAAATACAGGATGAAGGGAATTAAAAGATTTAATGCACATTTCTTCAAGGATAGTATTTCTGTATTGGCAAAATTTGAGAATAAATGGGTCTGGAACGAAAAAAAAAAAAAAA
